# A Comprehensive Proteogenomic and Spatial Analysis of Innate and Acquired Resistance of Metastatic Melanoma to Immune Checkpoint Blockade Therapies

**DOI:** 10.1101/2024.09.12.612675

**Authors:** Shiyou Wei, Kuang Du, Hongbin Lan, Zhenyu Yang, Yulan Deng, Zhi Wei, Dennie T. Frederick, Jinho Lee, Marilyne Labrie, Tian Tian, Tabea Moll, Yeqing Chen, Ryan J. Sullivan, Gordon Mills, Genevieve M. Boland, Keith T. Flaherty, Lunxu Liu, Meenhard Herlyn, Gao Zhang

## Abstract

While a subset of patients with metastatic melanoma achieves durable responses to immune checkpoint blockade (ICB) therapies, the majority ultimately exhibit either innate or acquired resistance to these treatments. However, the molecular mechanisms underlying resistance to ICB therapies remain elusive and are warranted to elucidate. Here, we comprehensively investigated the tumor and tumor immune microenvironment (TIME) of paired pre- and post-treatment tumor specimens from metastatic melanoma patients who were primary or secondary resistance to anti-CTLA-4 and/or anti-PD-1/PD-L1 therapies. Differentially expressed gene (DEG) analysis and single-sample gene set enrichment analysis (ssGSEA) with transcriptomic data identified cell cycle and c-MYC signaling as pathway-based resistance signatures. And weighted gene co-expression network analysis (WGCNA) revealed the activation of a cross-resistance meta-program involving key signaling pathways related to tumor progression in ICB resistant melanoma. Moreover, spatially-resolved, image-based immune monitoring analysis by using NanoString’s digital spatial profiling (DSP) and Cyclic Immunofluorescence (CyCIF) showed infiltration of suppressive immune cells in the tumor microenvironment of melanoma with resistance to ICB therapies. Our study reveals the molecular mechanisms underlying resistance to ICB therapies in patients with metastatic melanoma by conducting such integrated analyses of multi-dimensional data, and provides rationale for salvage therapies that will potentially overcome resistance to ICB therapies.

**Statement of translational relevance:** This study paves the way for the creation of innovative therapeutic strategies, aimed at subverting resistance to immune checkpoint blockade (ICB) therapies in metastatic melanoma patients. By unraveling the specific molecular mechanisms underlying resistance, scientists can design effective alternative treatments that target pathways such as pathways associated with cell cycle dysregulation and c-MYC signaling. Furthermore, through the application of advanced immune monitoring techniques such as NanoString Digital Spatial Profiling (DSP) and Cyclic Immunofluorescence (CyCIF), this study has significantly enriched our understanding of the tumor microenvironment. This enhanced characterization facilitates the discovery of potential biomarkers that may forecast a patient’s response to ICB treatment. Ultimately, these advancements could potentially refine patient outcomes and foster the development of more personalized cancer treatments in the future.

## Introduction

The advent of immune checkpoint blockade (ICB) therapies has significantly transformed the treatment landscape for metastatic melanoma, an aggressive and highly lethal form of skin cancer^1^. Although ICB therapies have markedly improved response rates and overall survival for some patients, a substantial proportion either exhibit innate resistance or develop acquired resistance, leading to treatment failure and disease progression^2–4^. Understanding the molecular mechanisms underlying both types of resistance is essential for optimizing therapeutic outcomes.

Innate resistance refers to a tumor’s intrinsic inability to respond to ICB therapies. Several molecular mechanisms contribute to this resistance. Tumors with a low tumor mutational burden (TMB) present fewer neoantigens, reducing immunogenicity and limiting the immune response to ICB therapies^5^. Defects in the antigen presentation machinery can impair immune cell recognition of cancer cells, leading to immune evasion^6^. This may occur through genetic alterations or epigenetic silencing of genes involved in antigen presentation, such as MHC class I, β2-microglobulin, and transporters associated with antigen processing^6^. Activation of alternative immune checkpoints, beyond the well-studied PD-1 and CTLA-4, can also suppress T cell function and contribute to innate resistance^6,7^. Additionally, the presence of immunosuppressive cells, including regulatory T cells (Tregs), myeloid-derived suppressor cells (MDSCs), and tumor-associated macrophages (TAMs), can dampen the anti-tumor immune response, resulting in resistance to ICB therapies^8^.

Acquired resistance occurs when a tumor initially responds to ICB therapy but eventually develops resistance. This resistance can be driven by several mechanisms. Immune selection pressure can lead to the emergence of tumor cell clones with reduced or lost expression of tumor antigens, preventing recognition by the immune system and facilitating immune escape^9^. Genetic alterations in key immune signaling pathways, such as the JAK/STAT and IFN-γ pathways, can also contribute to immune escape and acquired resistance^10^. Loss-of-function mutations in genes such as JAK1, JAK2, and the IFN-γ receptor can impair the tumor’s sensitivity to immune-mediated attack^10^. Under selective pressure, tumors may upregulate other immune checkpoints or negative regulators, such as TIM-3, LAG-3, VISTA, and IDO, to suppress T cell function and evade immune attack^11^. Additionally, dynamic changes in the tumor microenvironment (TME) during ICB therapy—including the recruitment of immunosuppressive cells, the development of an immunosuppressive extracellular matrix, and alterations in cytokine and chemokine expression—can modulate immune cell infiltration and function, contributing to acquired resistance^12^.

A comprehensive understanding of the molecular mechanisms underlying innate and acquired resistance to ICB therapies is crucial for developing novel therapeutic strategies and optimizing clinical outcomes in metastatic melanoma. To date, most studies have focused on analyzing patient-matched paired pre- and on-treatment tumor specimens to elucidate early adaptive responses to ICB therapies and identify predictive biomarkers^13–15^. Integrated analyses of transcriptomic and immune profiling data have revealed that activated T cell signatures and T cell populations are enriched in responders treated with ICB therapies^15^. Furthermore, transcriptomic data analysis from tumor specimens before and during anti-PD-1 therapy has shown increases in distinct immune cell subsets, activation of specific transcriptional networks, and upregulation of immune checkpoint genes in responders^16^. Studies using NanoString nCounter RNA data have also demonstrated that adaptive immune signatures are associated with response to ICB^17^.

In this study, we present a comprehensive analysis of RNA sequencing (RNAseq) data to reveal the transcriptomic landscape of innate and acquired resistance to ICB therapies in patients with metastatic melanoma. To our knowledge, this represents the largest cohort with RNAseq data available for patient-matched paired pre- and post-treatment tumor specimens from patients treated with ICB therapies. Our analysis provides a holistic understanding of the intricate interplay between cancer cells and the tumor microenvironment at the pathway level. By identifying key biomarkers, signaling pathways, and potential targets for therapeutic intervention, our integrative approach offers valuable insights into the molecular determinants of resistance. Furthermore, spatially-resolved, image-based immune monitoring analysis by using NanoString’s digital spatial profiling (DSP) and Cyclic Immunofluorescence (CyCIF) were performed to show the infiltration of immune cells in the tumor microenvironment of melanoma with resistance to ICB therapies. These findings lay the foundation for developing more effective, personalized treatment strategies for patients with metastatic melanoma.

## Results

### Patient Cohorts

In this study, we established a new cohort of patients with metastatic melanoma from Massachusetts General Hospital (MGH) and integrated this cohort with two published datasets^18–21^. RNA sequencing (RNAseq) data of patient-matched pre- and post-treatment tumor specimens were available for this integrated cohort at the bulk tumor level. “Post-treatment” specimens are defined as those from patients who exhibited either innate or acquired resistance to ICB therapies^22–24^. This cohort consisted of 80 tumor specimens derived from 25 patients treated with various ICB therapies, including anti-CTLA-4 monotherapy, anti-PD-1/PD-L1 monotherapy, anti-PD-1/PD-L1 monotherapy following anti-CTLA-4 monotherapy, and the combination of anti-PD-1 plus anti-CTLA-4 therapies, all of whom subsequently experienced disease progression (**Fig. 1a and Fig. 1b**).

**Figure 1.**
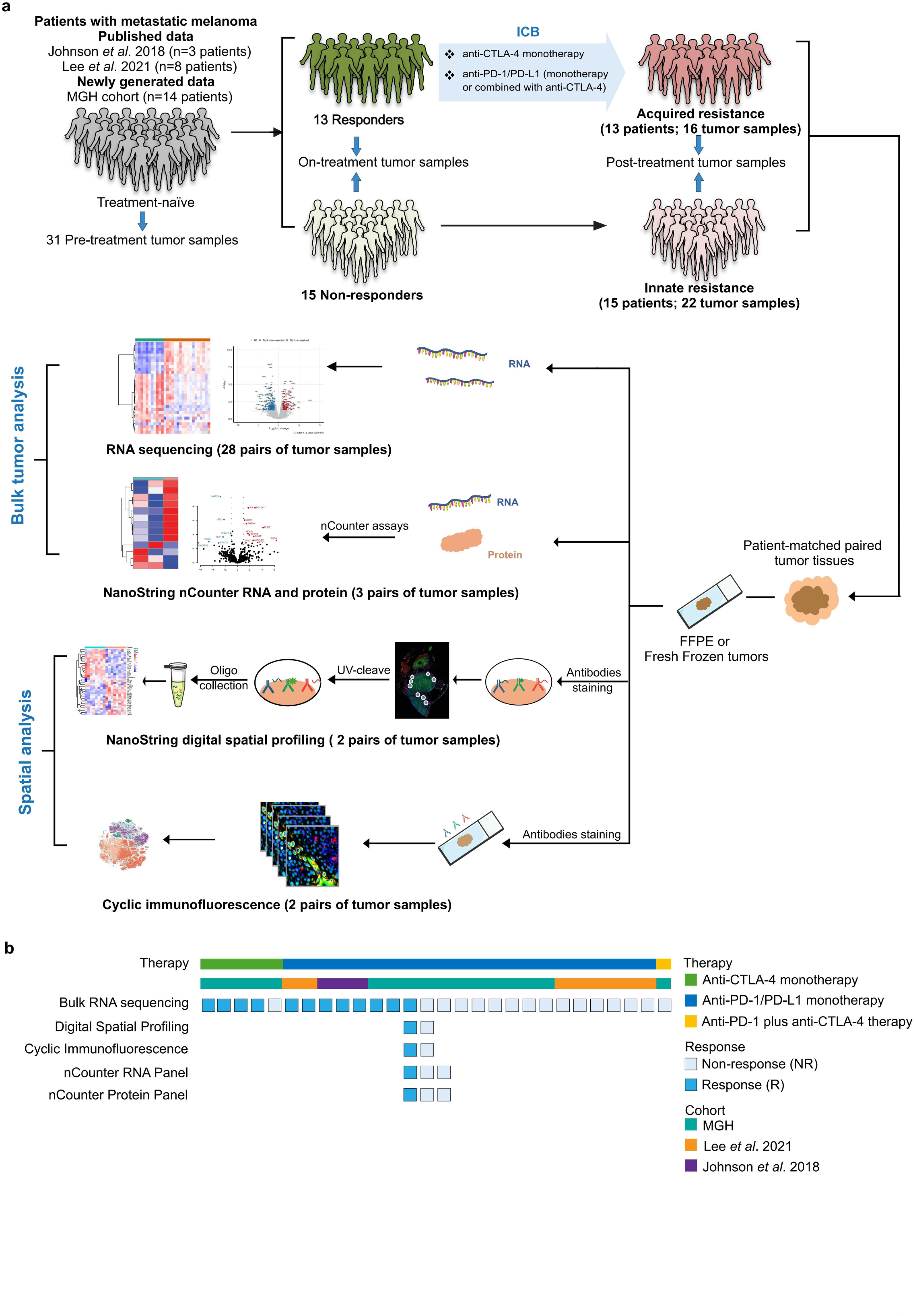
Patient Cohorts and Study Design. **(a)** Sample Collection and Analysis Workflow: This study incorporates data from patients with metastatic melanoma who received immune checkpoint blockade (ICB) therapies, combining cohorts from Johnson *et al.* (n=3), Lee *et al*. (n=8), and MGH (n=14). Paired pre-treatment, on-treatment, and/or post-treatment tumor specimens were collected from 13 responders and 15 non-responders and subjected to bulk tumor RNA sequencing (RNAseq). Both formalin-fixed paraffin-embedded (FFPE) and fresh frozen (FF) tumors were analyzed. In addition, four pairs of tumor samples underwent NanoString nCounter RNA and protein analysis, four pairs were analyzed using NanoString digital spatial profiling (DSP), and three pairs were subjected to Cyclic immunofluorescence (CycIF). The comprehensive experimental workflow is illustrated in this panel. **(b)** Analysis Approaches for Cohort Samples: This panel outlines the diverse analytical methodologies applied to samples from the Johnson *et al.,* Lee *et al.*, and MGH cohorts, emphasizing the different techniques used to investigate the molecular and spatial characteristics of the tumor specimens.

Specifically, three responders from the Johnson *et al*. study were included in this cohort, who received anti-PD-1 monotherapy and then progressed **(Fig. 1b and Table S1**)^18^. These patients had matched pre- and post-treatment tumor specimens’ transcriptomes sequenced, and their treatment responses were evaluated based on the Response Evaluation Criteria in Solid Tumors (RECIST). Additionally, from the Lee *et al*. study, two responders and six non-responders who received anti-PD-1 monotherapy were included in this cohort **(Fig. 1b and Table S1**)^19^. Tumor responses for this cohort were assessed using immune-related response criteria (irRC). For patients treated at MGH, seven responders and eight non-responders were included in this cohort, encompassing patients treated with anti-CTLA-4 monotherapy, anti-PD-1 monotherapy and the combination of anti-CTLA-4 plus anti-PD-1 therapies, as well as prior BRAF inhibitor (BRAFi) therapy **(Fig. 1b and Table S1**)^20,21^.

To serve as additional controls for tumor specimens at the time of acquired resistance to ICB therapies, matched pre- and on-treatment samples from 42 patients from the Riaz *et al*. study and 5 patients from MGH were also included^25^. These patients, classified as responders (R) or non-responders (NR) according to RECIST criteria, received anti-PD-1/PD-L1 therapy. To further explore molecular mechanisms underlying the development and evolution of acquired resistance to anti-PD-1 therapies, we included RNAseq data from two pairs of patient-matched pre-, on-, and post-treatment tumor specimens. Additionally, we extended our analysis to another published cohort of 5 patients with recurrent glioblastoma (rGBM) treated with anti-PD-1 therapy (Patients’ information please see **Table S1**)^26^.

To unravel cross-resistance mechanisms exhibited by patients with metastatic melanoma who were treated with molecularly targeted therapies or ICB therapies, we included two published cohorts comprising 25 patients who were initially treated with BRAFi and then subsequently relapsed^27,28^. Additionally, we included one patient from the MGH cohort treated with BRAFi, with RNAseq data available from pre- and post-treatment tumor specimens prior to receiving immunotherapy.

For two patients treated at MGH, we collected three pairs of tumor specimens subject to NanoString nCounter RNA and protein analysis. Additionally, two pairs of tumor specimens from these two patients were subject to NanoString digital spatial profiling (DSP) and cyclic immunofluorescence (CyCIF). These pairs included patient-matched samples at pre-treatment and at the time of innate or acquired resistance to anti-PD-1/anti-CTLA-4 therapy (**Fig.1b**).

In total, we assembled the largest cohort to date with integrated and multi-omics data generated, comprising 28 pairs of ICB-treated patient-matched pre- and post-treatment metastatic melanomas, 47 pairs of ICB-treated patient-matched pre- and on-treatment metastatic melanomas, 18 pairs of BRAFi-treated patient-matched pre- and post-treatment metastatic melanomas, 11 pairs of BRAFi-treated patient-matched pre- and on-treatment metastatic melanomas, and 5 pairs of ICB-treated patient-matched pre- and post-treatment GBM specimens. For all these tumor specimens, bulk tumor RNAseq data were available. Additionally, 3 pairs of ICB-treated patient-matched pre- and post-treatment metastatic melanomas were available for NanoString nCounter RNA and protein analysis, and two pairs were available for DSP and CyCIF analysis. This integrated and multi-omics data permitted us to carry out our subsequent computational analyses and investigations.

### The identification of Hallmark gene sets that are enriched in metastatic melanoma and GBM treated with molecularly targeted therapies and ICB therapies

To identify signaling pathways that were altered during and after anti-PD-1 therapy, we conducted single sample gene set enrichment analysis (ssGSEA) to derive scores for each Molecular Signatures Database (MSigDB) Hallmark gene set using leading-edge genes^29,30^. We compared ssGSEA scores of on-treatment tumor specimens to those of pre-treatment specimens for 19 patients with metastatic melanoma who initially responded to anti-PD-1 therapy (classified as responders, Rs). Similarly, we compared ssGSEA scores of post-treatment specimens to those of pre-treatment specimens for 10 patients with metastatic melanoma who initially responded but later acquired resistance to anti-PD-1 therapy (classified as non-responders, NRs).

The clustering analysis of these ssGSEA scores revealed a notable pattern in response to anti-PD-1 therapy. This pattern includes the dynamic changes of immune-related gene sets and tumor-related gene sets between initially responding to the ICB therapies and eventually developing acquired resistance (**Fig. 2a**). Specifically, a subset of immune-related gene sets was up-regulated in Rs’ on-treatment tumor specimens but down-regulated in NRs’ post-treatment specimens. These included *inflammatory response*, *interferon gamma response*, *interferon alpha response*, *IL6-JAK-STAT3 signaling*, *IL2-STAT5 signaling*, *allograft rejection* and *complement pathways*. Hereby we referred this subset as the immune meta-program.

**Figure 2.**
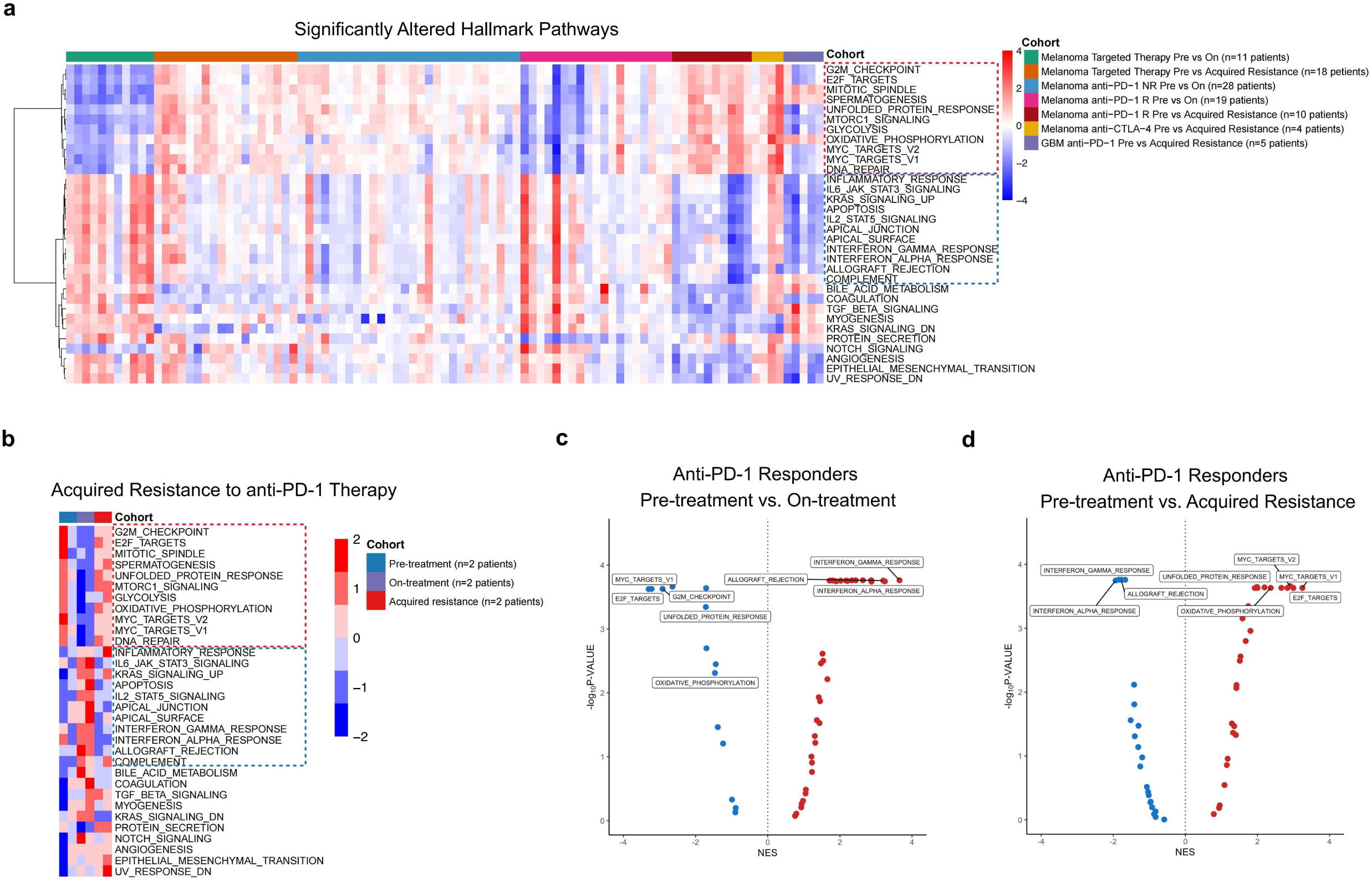
Significantly Enriched Hallmark Pathways Identified in On- or Post-treatment Tumor Specimens from Patients with Metastatic Melanoma or Glioblastoma Treated with ICB or Targeted Therapies Compared to Pre-treatment Specimens. (a) Heatmap of ssGSEA Scores: This heatmap presents the single-sample Gene Set Enrichment Analysis (ssGSEA) scores for significantly enriched Human MSigDB hallmark pathways, comparing tumor samples at the on-treatment or acquired resistance time points to matched pre-treatment samples. The samples originate from patients with metastatic melanoma treated with either targeted therapies or ICB therapies, including cohorts from MGH, Riaz *et al*., Hugo *et al*., Kwong *et al.,* and Zhao *et al*. (details provided in Fig. S1 and Table S1). Each column represents the difference in ssGSEA scores between a patient’s on- or post-treatment tumor sample and their pre-treatment sample, with each row corresponding to a significantly enriched Human MSigDB hallmark pathway. (b) Heatmap of ssGSEA Scores Over Time: This heatmap displays ssGSEA scores for significantly enriched Human MSigDB hallmark pathways in tumor samples at pre-treatment, on-treatment, and acquired resistance time points. These samples were obtained from two patients with acquired resistance to anti-PD-1 therapy from the MGH cohort, allowing for the observation of dynamic changes in pathway enrichment throughout the treatment timeline. (c) Volcano Plot of Pathway Regulation During Treatment: The volcano plot illustrates the down-regulated (blue) and up-regulated (red) Human MSigDB hallmark pathways by comparing tumor samples at the on-treatment time point to those at the pre-treatment time point. These samples were derived from anti-PD-1 responders from the MGH and Riaz et al. cohorts, highlighting significant changes in pathway activation or suppression in response to the treatment. (d) Volcano Plot of Pathway Regulation at Acquired Resistance: This volcano plot shows the down-regulated (blue) and up-regulated (red) Human MSigDB hallmark pathways by comparing tumor samples at the acquired resistance time point to those at the pre-treatment time point. The samples were derived from anti-PD-1 responders from the MGH, Johnson et al., and Lee et al. cohorts, emphasizing the pathways that become significantly altered as tumors develop resistance to the therapy.

Conversely, another subset of gene sets, including *cell cycle and DNA repair pathways* (G2M checkpoint, E2F targets, and mitotic spindle), *adaptive stress response pathways* (unfolded protein response (UPR) and oxidative phosphorylation (OXPHOS)), and *key tumor signaling pathways* (mTORC1 signaling, MYC targets V1 and V2), was down-regulated in Rs’ on-treatment specimens but up-regulated in NRs’ post-treatment specimens (**Fig. 2a**). Interestingly, we also observed the same pattern when comparing ssGSEA scores of tumor specimens exhibiting acquired resistance (n=18) to those of on-treatment specimens when patients with *BRAF*-mutant melanoma were treated with BRAFi (n=11) (**Fig. 2a**). We extended our analysis to three additional scenarios, including (1) acquired resistance to anti-CTLA-4 therapy, (2) on-treatment specimens from NRs treated with anti-PD-1 therapy and (3) in patients with recurrent glioblastoma (rGBM) who acquired resistance to anti-PD-1 therapy. In all scenarios, the same categories of MSigDB Hallmark gene sets were enriched in therapy-resistant tumors when compared to pre-treatment specimens, regardless of cancer type or therapy. Among them, cell cycle and DNA repair pathways, adaptive stress response pathways, and key tumor signaling pathways were included (**Fig. 2a**). This suggests that there is an existence of a common transcriptional program conferring cross-resistance to cancer therapies, referred as the cross-resistance meta-program.

Our further analysis of RNAseq data derived from two pairs of patient-matched pre-, on-, and post-anti-PD-1 treatment specimens, reinforced the dynamic nature of the cross-resistance meta-program. This meta-program was initially down-regulated upon anti-PD-1 treatment but gradually up-regulated in tumors that acquired resistance (**Fig. 2b**).

More specifically, three MSigDB Hallmark gene sets were up-regulated in on-treatment specimens from Rs treated with anti-PD-1 therapy, including *interferon alpha response*, *interferon gamma response* and *allograft rejection* (**Fig. 2c**). These gene sets were instead down-regulated in tumors derived from patients who acquired resistance to anti-PD-1 therapy (**Fig. 2d**). Similarly, five MSigDB Hallmark gene sets were down-regulated in on-treatment specimens from Rs, including *MYC targets V1*, *E2F targets*, *G2M checkpoint*, *UPR* and *OXPHOS* (**Fig. 2c**). All five of these gene sets were up-regulated in tumors from patients who acquired resistance to anti-PD-1 therapy except *G2M checkpoint*. Additionally, MYC targets V1 was up-regulated in resistant tumors (**Fig. 2d**).

In summary, our analysis revealed distinct expression patterns of two meta-programs when comparing tumor specimens that acquired resistance to anti-PD-1 therapy to those initially responding to the therapy. These findings provide valuable insights into molecular mechanisms underlying resistance to ICB therapies.

### Pathway-Based Signatures of Acquired Resistance to Anti-PD-1 Therapy in Metastatic Melanoma

In this study, we conducted a comprehensive analysis of gene expression profiles of tumor specimens collected from patients with metastatic melanoma before and after treatment with ICB therapies. Our objective was to identify differentially expressed genes (DEGs) and to identify molecular pathways associated with both acquired and innate resistance to anti-PD-1 and anti-CTLA-4 therapies. After applying stringent criteria for DEG identification and using |log2FC| ≥ 1and p<0.05, we revealed significant gene expression changes across different resistance scenarios. Specifically, for innate resistance to anti-CTLA-4 therapy, 582 DEGs were identified, including 423 up-regulated and 159 down-regulated genes (**Fig. S3a**). In the context of acquired resistance to anti-CTLA-4 therapy, we found 514 DEGs, comprising 172 up-regulated and 342 down-regulated genes (**Fig. S3b**). For tumors with innate resistance to anti-PD-1 therapy, there were 445 DEGs, with 321 genes up-regulated and 124 down-regulated (**Fig. S3c**). For tumors exhibiting acquired resistance to anti-PD-1 therapy, we identified 639 DEGs, including 226 up-regulated and 413 down-regulated genes compared to pre-treatment specimens (**Fig. S3d**).

To elucidate molecular mechanisms driving resistance to ICB therapies, we performed gene set enrichment analysis (GSEA) using the Chemical and Genetic Perturbations (CGP) collection from the MSigDB^30,31^. We focused on identifying pathway-based transcriptional signatures associated with innate and acquired resistance to anti-CTLA-4 and anti-PD-1 monotherapies. Specifically, we prioritized the top 10 up-regulated and down-regulated CGP pathways in post-treatment tumor specimens compared to their pre-treatment counterparts. Our analysis revealed several key insights into molecular pathways associated with resistance. Tumors with acquired resistance to anti-CTLA-4 therapy displayed the enrichment of pathways associated with cell cycle and proliferation, stem cell signaling, tumor metastasis, and tumor marker genes (**Fig. 3a and Fig. 3b**). In case of innate resistance to anti-PD-1 therapy, the enriched pathways included those involved in cell cycle regulation, tumor marker genes, and drug response mechanisms (**Fig. 3c and Fig. 3d**). Tumor specimens with acquired resistance to anti-PD-1 therapy showed the enrichment of pathways related to cell cycle regulation and proliferation, stem cell signaling, tumor invasiveness and metastasis, c-MYC signaling, and tumor marker genes. Additionally, these specimens exhibited gene signatures indicative of poor survival outcomes (**Fig. 3e and Fig. 3f**). Taken together, these findings highlight the complexity of molecular mechanisms of resistance exhibited by metastatic melanoma under therapeutic pressure and underscore the involvement of signaling pathways that are critical for cell cycle control, stem cell signaling, and tumor metastasis. The enrichment of c-MYC signaling and tumor marker genes further suggests that these pathways play a significant role in the development of resistance to ICB therapies.

**Figure 3.**
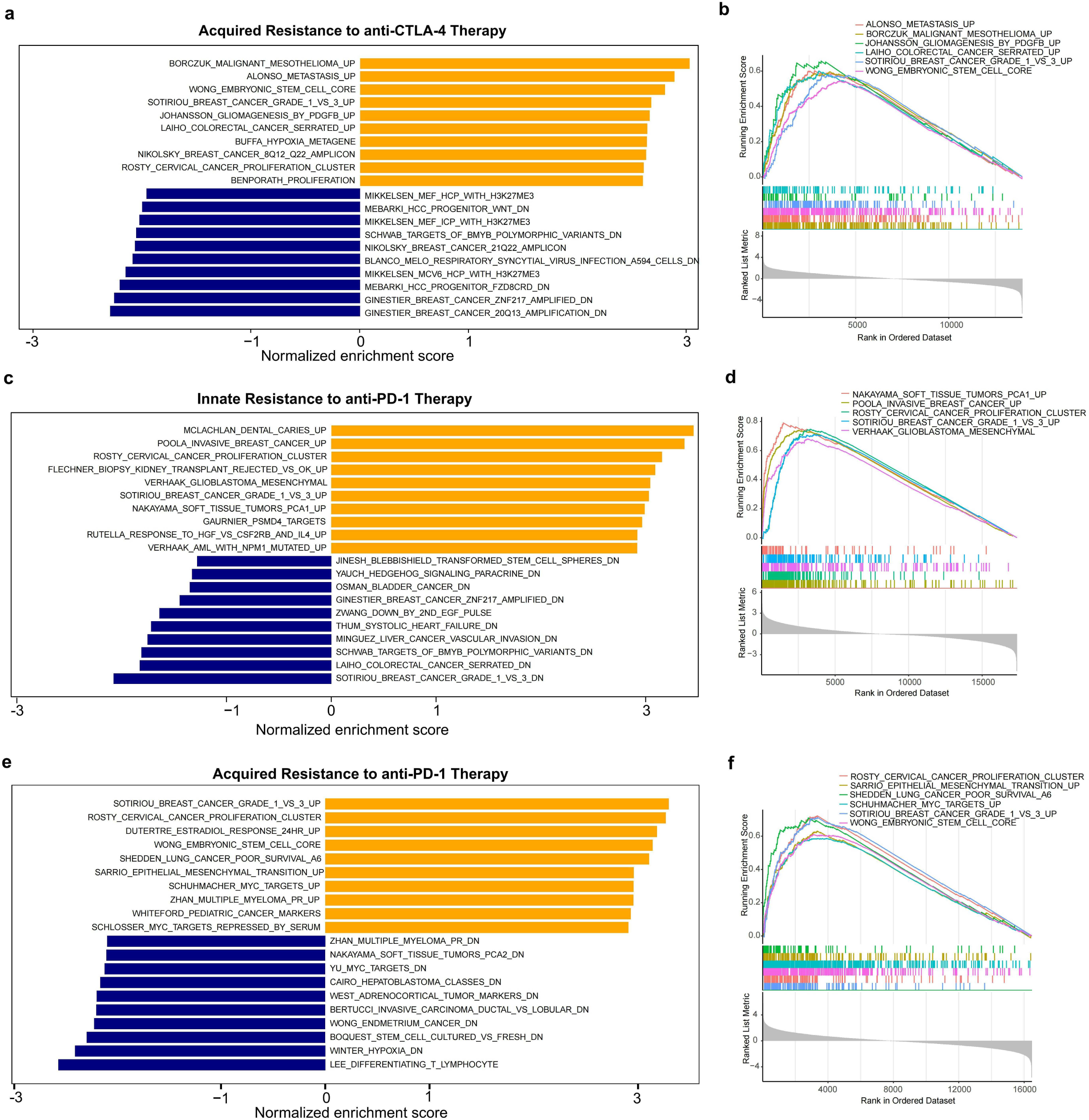
Pathway-Based Signature of Acquired or Innate Resistance to Anti-PD-1/Anti-CTLA-4 Therapy. (a) Two-Sided Bar Plot of CGP Pathways for Acquired Resistance to Anti-CTLA-4 Therapy: This bar plot shows absolute normalized enrichment scores for the top 10 up-regulated and down-regulated CGP pathways that were significantly different between tumor samples with acquired resistance to anti-CTLA-4 therapy and pre-treatment tumor specimens (FDR < 0.05). (b) Enrichment Plot for Acquired Resistance to Anti-CTLA-4 Therapy: This plot presents enrichment scores for selected CGP pathways that are up-regulated in tumor specimens with acquired resistance to anti-CTLA-4 therapy compared to pre-treatment tumor specimens. (c) Two-Sided Bar Plot of CGP Pathways for Innate Resistance to Anti-PD-1 Therapy: This bar plot shows absolute normalized enrichment scores for the top 10 up-regulated and down-regulated CGP pathways that were significantly different between tumor specimens with innate resistance to anti-PD-1 therapy and pre-treatment tumor specimens (FDR < 0.05). (d) Enrichment Plot for Innate Resistance to Anti-PD-1 Therapy: This plot displays enrichment scores for selected CGP pathways that are up-regulated in tumor specimens with innate resistance to anti-PD-1 therapy compared to pre-treatment tumor specimens. (e) Two-Sided Bar Plot of CGP Pathways for Acquired Resistance to Anti-PD-1 Therapy: This bar plot shows absolute normalized enrichment scores for the top 10 up-regulated and down-regulated CGP pathways that were significantly different between tumor specimens with acquired resistance to anti-PD-1 therapy and pre-treatment tumor specimens (FDR < 0.05). (f) Enrichment Plot for Acquired Resistance to Anti-PD-1 Therapy: This plot presents enrichment scores for selected CGP pathways that are up-regulated in tumor specimens with acquired resistance to anti-PD-1 therapy compared to pre-treatment tumor specimens.

In summary, our study provides a detailed molecular landscape of resistance to ICB therapies exhibited by metastatic melanoma. Pathway-based signatures that we identified offer valuable insights into the underlying mechanisms of therapeutic resistance and present potential targets for overcoming resistance. These findings could inform the development of more effective combination therapies and improve clinical outcomes for patients with metastatic melanoma.

### WGCNA Modules Conferring Resistance to Immune Checkpoint Blockade Therapies

To identify clusters of co-expressed genes associated with resistance to ICB therapies, we employed Weighted Gene Co-expression Network Analysis (WGCNA)^32^. We focused on uncovering molecular underpinnings of acquired resistance to anti-CTLA-4 therapy and both innate and acquired resistance to anti-PD-1 therapies. It is noteworthy that we generated networks based on gene expression data from various sample groups. Specifically, we included patient-matched tumor specimens at time points of pre-treatment and post-treatment (acquired resistance) to anti-CTLA-4 therapy from 4 patients with metastatic melanoma. We also included patient-matched samples at time points of pre-treatment and post-treatment (innate resistance) to anti-PD-1 therapy from 14 patients, among whom multiple post-treatment samples from 3 patients were included. Additionally, we included patient-matched tumor samples at time points of pre-treatment and post-treatment (acquired resistance) to anti-PD-1 therapy from 10 patients, including multiple post-treatment samples from 1 patient. Furthermore, we included patient-matched tumor samples from 18 patients with acquired resistance to BRAFi monotherapy, with multiple post-treatment samples from 9 patients (**Table S1**).

By setting the soft threshold power (β) at 10, we constructed a scale-free network and achieved an independence degree of 0.8 (**Fig. 4a**). This enabled us to cluster gene modules with similar expression patterns and determine module eigengene values to evaluate the overall expression patterns within these modules.

**Figure 4.**
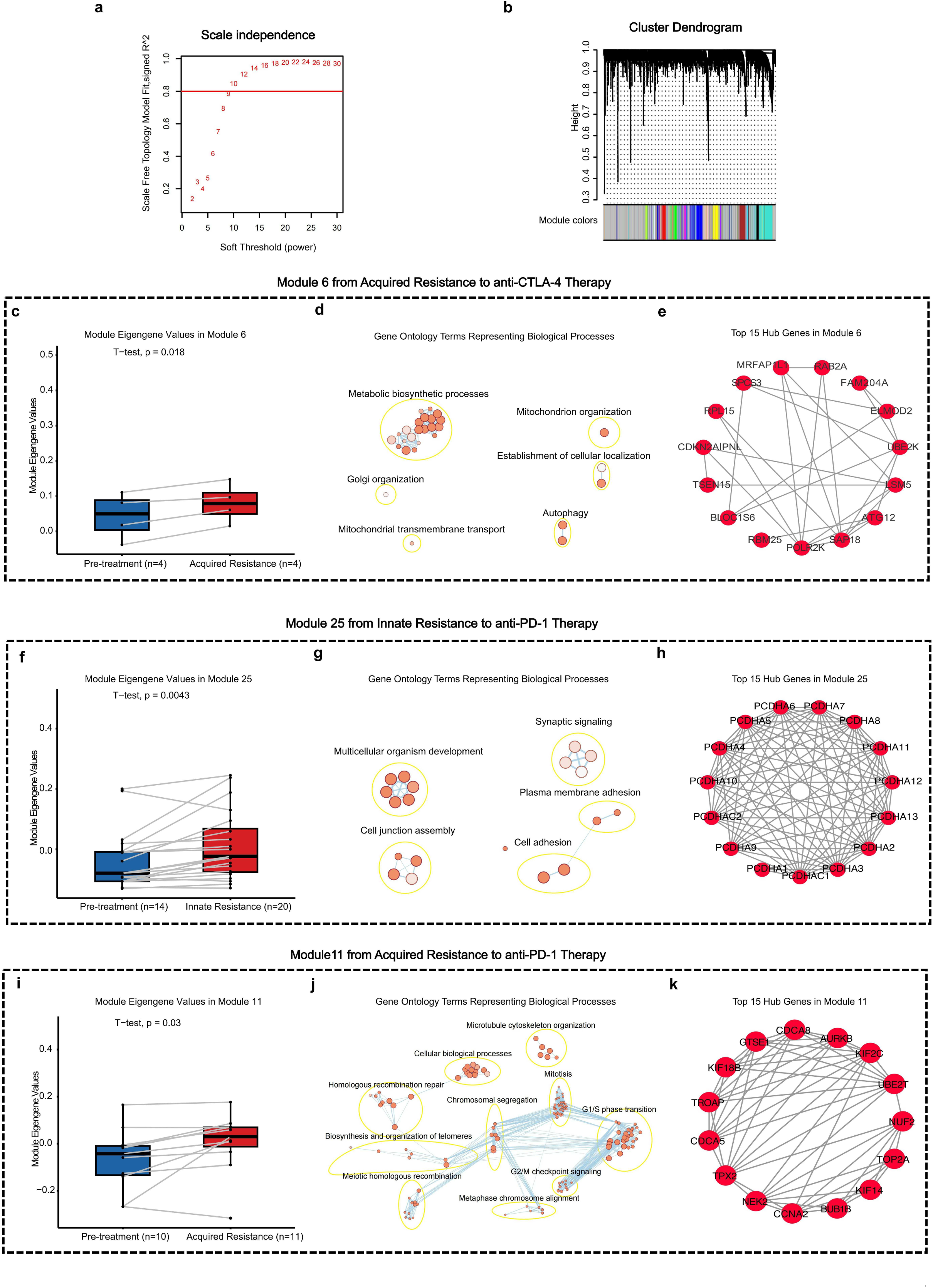
Modules Identified by WGCNA Conferring Resistance to Immune Checkpoint Blockade Therapies. (a) Scale Independence for Module Identification: This graph shows the scale independence achieved to identify modules. (b) Cluster Dendrogram: This dendrogram, created using hierarchical clustering, identifies modules based on gene expression data. (c) Boxplots of Module 6 Eigengene Values: These boxplots illustrate the module eigengene values for module 6, which is associated with acquired resistance to anti-CTLA-4 therapy. (d) Gene Ontology Annotation of Module 6: This annotation details the gene ontology terms for biological processes identified in module 6 by WGCNA. (e) Visualization of Top 15 Gene Hubs in Module 6: This visualization highlights the top 15 gene hubs within module 6. (f) Boxplots of Module 25 Eigengene Values: These boxplots show the module eigengene values for module 25, which is associated with innate resistance to anti-PD-1 therapy. (g) Gene Ontology Annotation of Module 25: This annotation details the gene ontology terms for biological processes identified in module 25 by WGCNA. (h) Visualization of Top 15 Gene Hubs in Module 25: This visualization presents the top 15 gene hubs within module 25. (i) Boxplots of Module 11 Eigengene Values: These boxplots display the module eigengene values for module 11, which is linked to acquired resistance to anti-PD-1 therapy. (j) Gene Ontology Annotation of Module 11: This annotation provides the gene ontology terms for biological processes identified in module 11 by WGCNA. (k) Visualization of Top 15 Gene Hubs in Module 11: This visualization showcases the top 15 gene hubs within module 11.

Our WGCNA analysis identified 4 significant co-expression modules, including module 10 (ME10), module 6 (ME6), module 25 (ME25), and module 11 (ME11) (**Fig. 4c**, **Fig. 4f**, **Fig. 4i and Fig. S4a**). ME10, associated with acquired resistance to BRAFi, showed enrichment in biological processes related to multicellular reproduction, as revealed by Gene Ontology (GO) enrichment analysis (**Fig. S4b**). ME6, associated with acquired resistance to anti-CTLA-4 therapy, was predominantly correlated with biosynthesis and metabolism processes (**Fig. 4d**). ME25, linked to innate resistance to anti-PD-1 therapy, was associated with pathways related to multicellular development processes and cell junction assembly (**Fig. 4g**). ME11, related to acquired resistance to anti-PD-1 therapy, exhibited a high correlation with biological processes related to cell cycle and DNA repair, including microtubule cytoskeleton organization, mitotic cell cycle processes, metaphase chromosome alignment, G1/S cell cycle phases, G2/M checkpoint signaling, chromosome segregation, meiotic homologous recombination, and homologous recombinational repair (**Fig. 4d**). This finding is well aligned with the result of our MSigDB Hallmark gene set analysis, which indicated that cell cycle and DNA repair pathways are potentially related to acquired resistance to ICB therapies.

Further analysis of hub genes within these modules allowed us to identify their associations with various types of therapeutic resistance. We constructed co-expression networks and selected pairs with a weight greater than 0.05, leading to the identification of the top 20 ranked significant gene hubs in ME10 (**Fig. S4c**) and top 15 ranked significant gene hubs in ME6, ME25 and ME11(**Fig. 4e**, **Fig. 4h, and Fig. 4k**). Of the 20 genes in ME10, TMIE, CDKL4, SCN11A, MYT1L, JPH3, SYT13 and NPFFR2 genes associated with neurological processes; ARHGAP15, NLRP4, ITGA2B and CDH20 genes associated with immune response, and SPAG6, TEX19 genes associated with reproduction process were found to be markedly enriched in BRAFi resistance. (**Fig. S4c**). ME11, associated with acquired resistance to anti-PD-1 therapy, contained gene hubs such as CDCA8, AURKB, KIF2C, KIF14, KIF18B, BUB1B, CCNA2, NEK2, TPX2, CDCA5, and NUF2, which are primarily associated with cell cycle regulation, division, and mitosis, while genes like UBE2T, GTSE1, and TOP2A are related to DNA repair processes (**Fig. 4e**). For ME25, gene hubs including PCDHA 1-13, PCDHAC1, and PCDHAC2 were identified; these genes are potentially involved in establishing and maintaining specific neuronal connections and are strongly associated with cell adhesion (**Fig. 4h**). In ME6, genes such as POLR2K, SPCS3, UBE2K, RAB2A, RPL15, and ATG12 were identified, all of which are related to biosynthesis and metabolism (**Fig. 4k**).

Overall, our WGCNA analysis revealed four distinct co-expression modules and hub genes associated with them. These modules are linked with acquired resistance to BRAFi targeted therapy, acquired resistance to anti-CTLA-4 therapy, innate resistance to anti-PD-1 therapy, and acquired resistance to anti-PD-1 therapy. These findings provide valuable insights into molecular mechanisms underlying therapeutic resistance and suggest potential targets for overcoming resistance in metastatic melanoma.

### Integrated Analyses of NanoString nCounter RNA and Protein Data

To analyze alterations in cancer-related signaling pathways by immune checkpoint blockade (ICB) therapies exhibited by post-treatment tumor specimens as compared to pre-treatment specimens, we further utilized the RNA Pan-Cancer Pathways Panel to assess mRNA transcripts and the Protein Solid Tumor Panel to examine 13 phosphorylated proteins.

For MGH patient 208 with innate resistance to anti-CTLA-4 therapy, our results indicated there was an up-regulation of the MAPK-RAS and PI3K signaling pathways in post-treatment tumor specimens as compared to pre-treatment tumors, while the proliferation pathway was down-regulated (**Fig. 5a**). At the phosphorylated protein level, we observed increases in p-c-Raf^Ser259^, p-MEK1/2^Ser217/221^, and p-ERK1/2^Thr202/Tyr204^, all of which were involved in the MAPK-RAS signaling pathway (**Fig. 5b**). Additionally, we observed increases in p-PRAS40^Thr246^ and p-S6^Ser235/236^ which were key players of the mTOR signaling pathway (**Fig. 5b**). Moreover, proteins such as p-GSK3B^Ser9^ and p-Akt^Ser473^ were up-regulated in the PI3K/Akt signaling pathway. Conversely, the down-regulation of p-Tuberin^Thr1462^, a key player in the mTOR pathway, and p-Histone H3^Ser10^, crucially linked to proliferation pathways, occurred (**Fig. 5b**).

**Figure 5.**
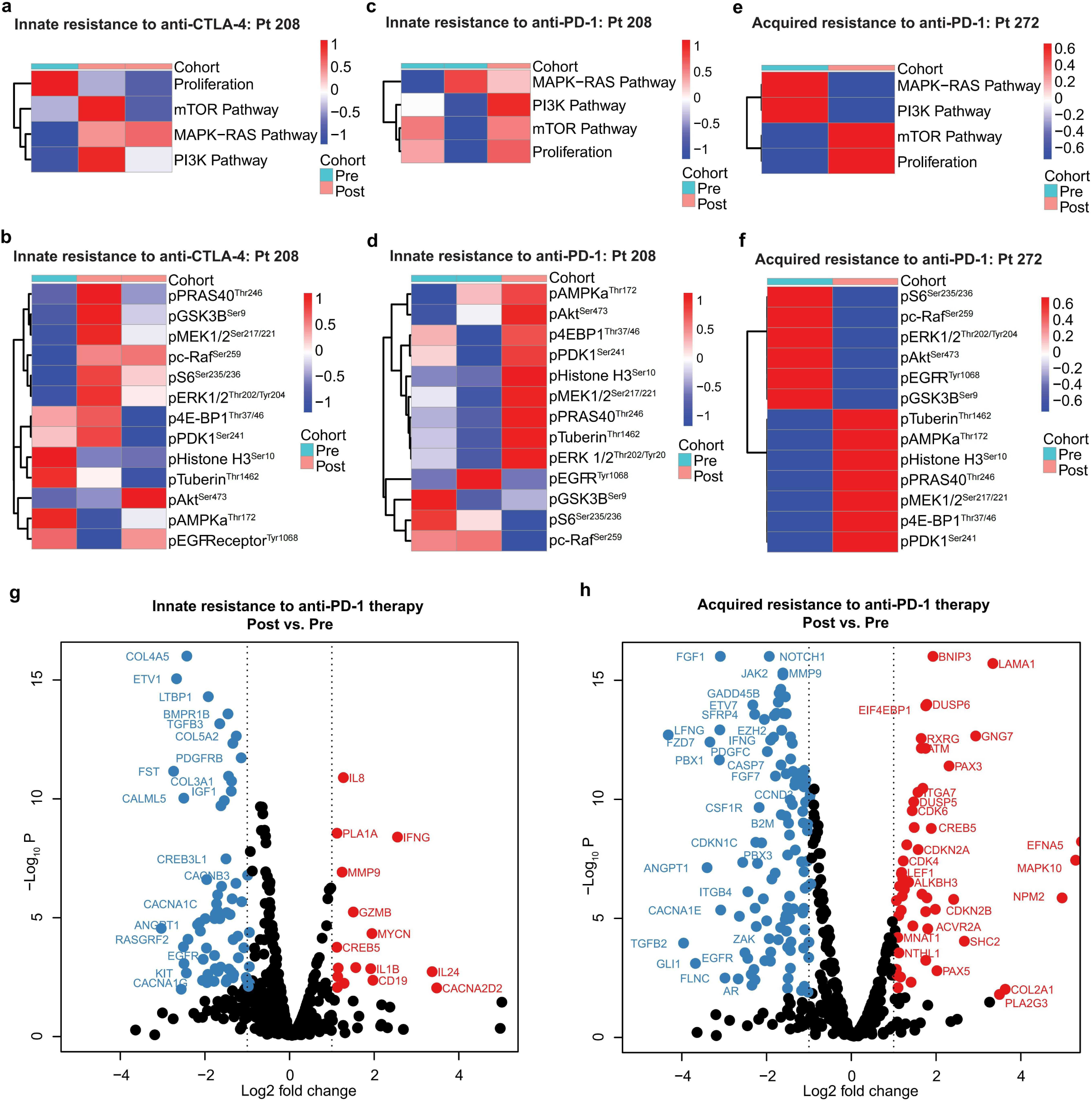
Integrated Analyses of NanoString nCounter RNA and Protein Data. Geomean normalized nCounter data was used for the pathway analysis and differential expression analysis. Pathway scores were weighted and calculated based on target specificity in each signaling pathway. (a) Heatmap showing four pathways for MGH patient 208 with innate resistance to anti-CTLA-4 therapy analyzed by NanoString nCounter protein platform. (b) Heatmap showing 13 phosphorylated proteins for MGH patient 208 with innate resistance to anti-CTLA-4 therapy analyzed by NanoString nCounter protein platform. (c) Heatmap showing four pathways for MGH patient 208 with innate resistance to anti-PD-1 therapy analyzed by NanoString nCounter protein platform. (d) Heatmap showing 13 phosphorylated proteins for MGH patient 208 with innate resistance to anti-PD-1 therapy analyzed by NanoString nCounter protein platform. (e) Heatmap showing four pathways for MGH Patient 272 with acquired resistance to anti-PD-1 therapy analyzed by NanoString nCounter protein platform. (f) Heatmap showing 13 phosphorylated proteins for MGH Patient 272 with acquired resistance to anti-PD-1 therapy analyzed by NanoString nCounter protein platform. (g) Volcano plot showing differential mRNA transcripts for post-treatment tumor specimens with innate resistance to anti-PD-1 therapy compared to pre-treatment tumor specimens analyzed by NanoString nCounter RNA platform. (h) Volcano plot showing differential mRNA transcripts for post-treatment tumor specimens of patients with acquired resistance to anti-PD-1 therapy compared to pre-treatment tumor specimens analyzed by NanoString nCounter RNA platform.

For MGH patient 208 with innate resistance to anti-PD-1 therapy, our results revealed that PI3K and proliferation pathways were upregulated (**Fig. 5c**). Moreover, phosphorylated proteins included p-Akt^Ser473^, p-4EBP1^Thr37/46^, p-PDK1^Ser241^, and p-PRAS40^Thr246^ which are associated with the PI3K pathway, along with p-Histone H3^Ser10^, involved in the proliferation pathway, were found to be upregulated in the post-treatment tumor specimens in comparison to the pre-treatment tumors (**Fig. 5d**).

In the case of MGH patient 272 with acquired resistance to anti-PD-1 therapy, the MAPK-RAS and PI3K signaling pathways were down-regulated but the mTOR and proliferation pathways were up-regulated in post-treatment tumor samples as compared to pre-treatment samples (**Fig. 5e**). The down-regulated phosphorylated proteins included p-S6^Ser235/236^, p-c-Raf^Ser259^, p-ERK1/2^Thr202/Tyr204^, p-Akt^Ser473^, p-EGFR^Tyr1068^ and p-GSK3B^Ser9^. Conversely, the up-regulated phosphorylated proteins included p-Tuberin^Thr1462^, p-AMPKa^Thr172^, p-Histone H3^Ser10^, p-PRAS40^Thr246^, p-MEK1/2^Ser217/221^, p-4E-BP1^Thr37/46^ and p-PDK1^Ser241^(**Fig. 5f**).

nCounter RNA analysis compared post-treatment tumor specimens to pre-treatment tumor specimens from patients with metastatic melanoma who underwent anti-PD-1 therapy |log2FC| ≥1and p<0.05. In tumor specimens demonstrating innate resistance to anti-PD-1 therapy, we observed up-regulated gene expressions such as IL8, IL1B, IFNG, and IL24, associated with inflammatory response modulation (**Fig. 5g**). Additionally, MYCN and CREB5, involved in transcriptional regulation were also upregulated. Moreover, CACNA2D2, related to calcium signaling and channels, and PLA1A, MMP9, and GZMB, which are related to pathways including lipid metabolism, extracellular matrix remodeling and degradation, and apoptosis, were found to be upregulated. (**Fig. 5g**). In contrast, we observed down-regulated mRNA transcripts including TGFB3, PDGFRB, FST, IGF1, ANGPT1, RASGRF2, and KIT, which contributes to the dysregulation of immune responses (**Fig. 5g and Table S6**).

In tumor specimens demonstrating acquired resistance to anti-PD-1 therapy, we identified several key gene expressions upregulated compared to pre-treatment tumors. Specifically, RAX3, functioning as a transcription factor, along with LAMA1, PLA2G3, and COL2A1, linked to extracellular matrix components, as well as MAPK10 and EFNA5, pivotal in cellular processes and signaling, showed increased expression levels in these resistant tumor specimens (**Fig. 5h**). Conversely, significant alterations were observed in the gene expression of key players within various signaling pathways. For instance, TGFB2, a vital component of the TGF-beta pathway, were down-regulated. Similarly, NOTCH1, a crucial contributor to the notch signaling pathway, and JAK2, a pivotal member of the JAK-STAT pathway, also down-regulated in expression levels. Moreover, down-regulated gene expressions were identified in genes associated with cell cycle regulation and cell proliferation, exemplified by the decreased expression of CDKN1C and CCND2 (**Fig.5h & Table S7**).

Taken together, our data revealed that the PI3K signaling pathway was up-regulated in innate resistance to anti-PD-1 and anti-CTLA-4 therapies, while it was down-regulated in acquiring resistance to anti-PD-1 therapy. Additionally, the down-regulation of MAPK-RAS and PI3K signaling pathways and up-regulation of the mTOR and proliferation pathways may confer acquired resistance to anti-PD-1 therapy. Our findings showed different levels of phosphorylated proteins and gene expressions that demonstrated signaling pathway alterations, offering valuable insights into the mechanisms underlying resistance to ICB therapies.

### Spatially Resolved Immune Profiling of ICB-resistant Metastatic Melanomas Using NanoString DSP Platform

To gain a deeper insight into dynamics that orchestrated immune response of metastatic melanoma to ICB therapies, we employed NanoString GeoMx® Digital Spatial Profiling (DSP) technology^33^. Particularly, we assembled a panel of 33 markers of immuno-oncology, which were meticulously assessed within each ROI, providing valuable insights into the dynamics of the immune response (**Fig. 6a**). In these experiments, we selected two pairs of patient-matched pre- and post-treatment FFPE tumor specimens as representative examples, including one pair of innate resistance to anti-PD-1 therapy and one pair of acquired resistance to anti-PD-1 therapy. Our investigation focused on tumor regions with positive staining for the melanocytic marker S100, which also exhibited a high content of immune cells as determined by CD45 positivity. These criteria allowed us to selectively choose regions of interest (ROIs) within the tumor. We selected an average of 8 ROIs from each formalin-fixed paraffin-embedded (FFPE) tumor specimen with examples shown in **Fig. 6b** and **Fig. 6c**.

**Figure 6.**
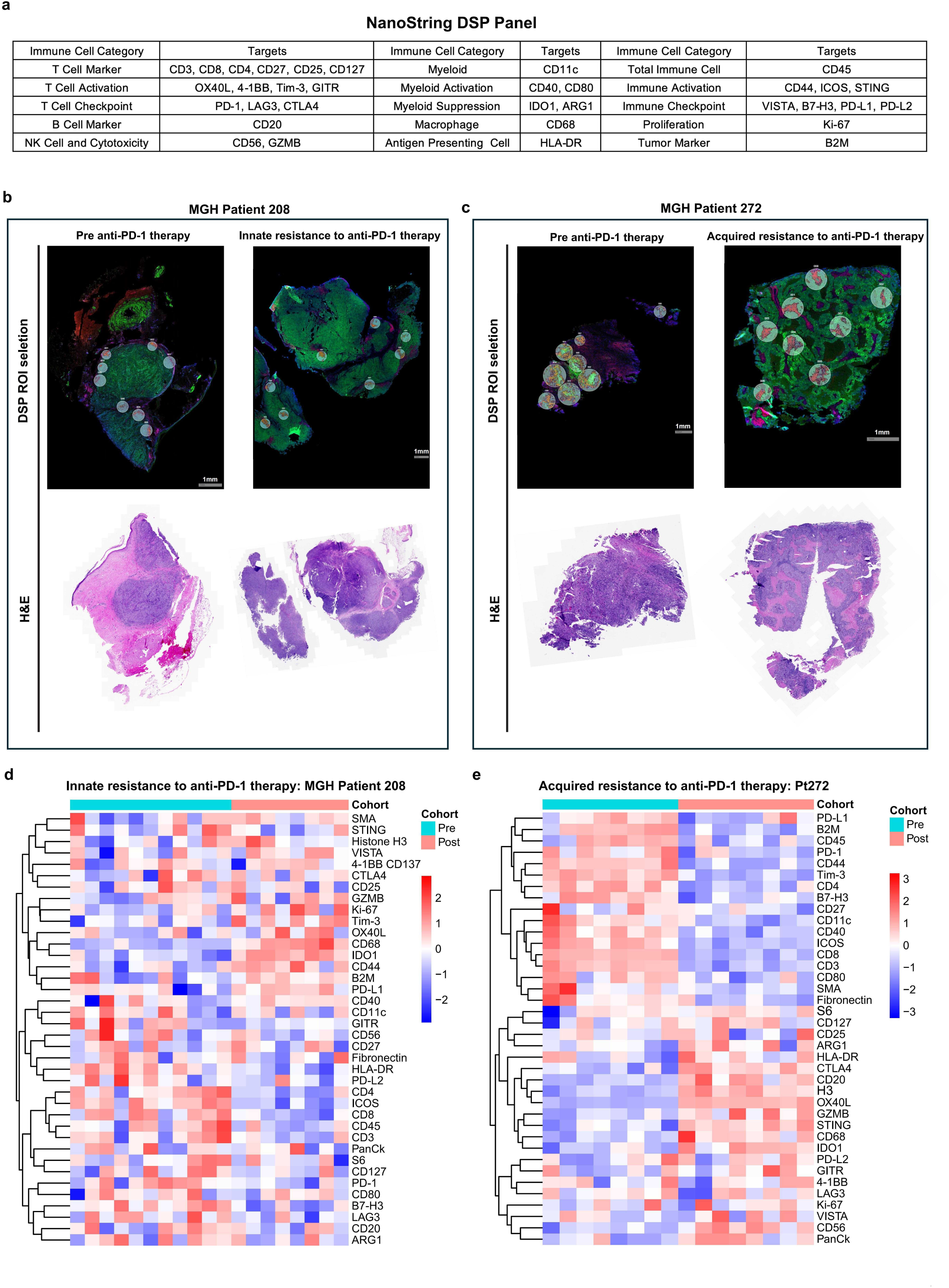
Spatially Resolved Immune Profiling by NanoString DSP Platform. DSP 33 immune markers were analyzed in CD45 positive regions for immune profiling. (a) NanoString DSP Immune Panel of 33 Proteins: This panel illustrates the comprehensive array of 33 immune-related proteins analyzed using the NanoString DSP technology. These proteins were selected for their critical roles in immune signaling pathways and tumor-immune interactions. (b) Representative Figures for DSP Region of Interest (ROI) Selection and H&E Staining of Tumor Specimens of MGH patient 208: This figure showcases the regions of interest (ROIs) selected for DSP analysis and H&E-stained sections from tumor specimens of MGH patient 208 at two critical time points: pre-treatment and post-treatment with anti-PD-1 therapy, where the latter represents the state of innate resistance. (c) Representative Figures for DSP ROIs Selection and H&E Staining of Tumor Specimens of MGH Patient 272: This figure showcases the ROIs selected for DSP analysis and H&E-stained sections from tumor specimens of MGH patient 272. The figures capture these regions at crucial time points: pre-anti-PD-1 therapy and acquired resistance to anti-PD-1 therapy. (d) DSP Protein Expression Heatmap for Innate Resistance to Anti-PD-1 Therapy in MGH Patient 208: This heatmap displays the protein expression data obtained from 11 ROIs in pre-treatment tumor specimens and 8 ROIs in post-treatment tumor specimens, illustrating innate resistance to anti-PD-1 therapy. The figure highlights key changes in immune protein expression associated with the innate resistance mechanism. (e) DSP Protein Expression Heatmap for Acquired Resistance to Anti-PD-1 Therapy in MGH Patient 272: This heatmap provides a detailed comparison of immune protein expression between 8 ROIs from pre-treatment tumor specimens and 8 ROIs from post-treatment tumor specimens, representing the state of acquired resistance to anti-PD-1 therapy in MGH patient 272. The heatmap elucidates significant alterations in immune protein profiles linked to the acquired resistance.

For tumor specimens exhibiting innate resistance to anti-PD-1 therapy, the analysis demonstrated a reduction in immunosuppression in post-treatment tumor samples as compared to pre-treatment tumor samples, as evidenced by the down-regulation of immune checkpoints such as PD-L2, LAG3, B7-H3 and T cell checkpoints such as PD-1 (**Fig. 6d**). Furthermore, there is an observed increase in T cell activation, immune activation markers, NK cell markers, and myeloid activation markers, which include STING, 4-1BB, GZMB, Tim-3, OX40L, CD44, and CD40, potentially enhancing the immune response (**Fig. 6d**). However, these samples also displayed an up-regulation of tumor markers and proliferation markers, such as Ki67, potentially indicative of tumor progression (**Fig. 6d**). Additionally, down-regulation of HLA-DR indicates a decrease in antigen-presenting cells, which may contribute to innate resistance to anti-PD-1 therapy (**Fig. 6d**).

For tumor specimens displaying acquired resistance to anti-PD-1 therapy, the comparison between pre- and post-treatment tumor samples revealed a decrease in the recruitment and activation of overall immune and myeloid cells, as indicated by the down-regulation of CD45, ICOS, CD11c, CD40 and CD80 (**Fig. 6e**). Despite of the up-regulation of markers of T cells, B cells, NK cells, antigen-presenting cells and immune activation, which typically suggest there is a positive immune response conducive to tumor eradication, those post-treatment samples also demonstrated there was an up-regulation of IDO1, ARG1, CTLA4 and LAG3. These markers are associated with myeloid suppression and T cell checkpoint activity, suggesting that an increase in immunosuppressive activity might contribute to acquired resistance to anti-PD-1 therapy. Furthermore, the heightened expression of Ki-67, a marker indicative of cell proliferation, could suggest tumor growth persisting despite anti-PD-1 therapy (**Fig. 6e**). The complex interplay of these up-regulated and down-regulated markers provides valuable insights into the intricate immune dynamics and the significant role of myeloid suppression in promoting acquired resistance to anti-PD-1 therapy.

Overall, the DSP results revealed a decrease in immunosuppression and an increase in immune cells and activation, while the down-regulation of antigen-presenting cells may play a crucial role in the innate resistance to anti-PD-1 therapy. Furthermore, despite of the up-regulation of immune cells and their activation, we noted a simultaneous up-regulation of myeloid suppression and T cell checkpoint activities, which may contribute to acquired resistance to anti-PD-1 therapy. Taken together, our results highlighted the complex interplay among different immune components within the tumor immune microenvironment that influences response of metastatic melanoma to ICB therapies. Understanding these mechanisms is crucial for developing strategies to overcome resistance and to improve the efficacy of immune-based cancer treatments.

### Cyclic Immunofluorescence (CyCIF) Identifies Dynamic Changes in Cell Types and States

To further elucidate molecular mechanisms at the spatial level underlying innate or acquired resistance of metastatic melanoma to immune checkpoint blockade (ICB) therapies, we conducted a spatially-oriented single-cell proteomics analysis by performing cyclic immunofluorescence (CyCIF) staining of formalin-fixed paraffin-embedded (FFPE) tissue slides from two patients, Pt 208, and Pt 272^34^. For this spatial analysis at the single cell level encompassing 23 proteins, we identified cell subtypes (S100^+^ melanoma cells and CD45^+^ immune cells) and characterized cell states, including epithelial-mesenchymal transition, transcriptional repression, DNA damage response (DDR), cell cycle/proliferation, and key signaling pathways (RTK, MAPK, and PI3K-AKT-mTOR signaling axes) (**Fig. 7a**).

**Figure 7.**
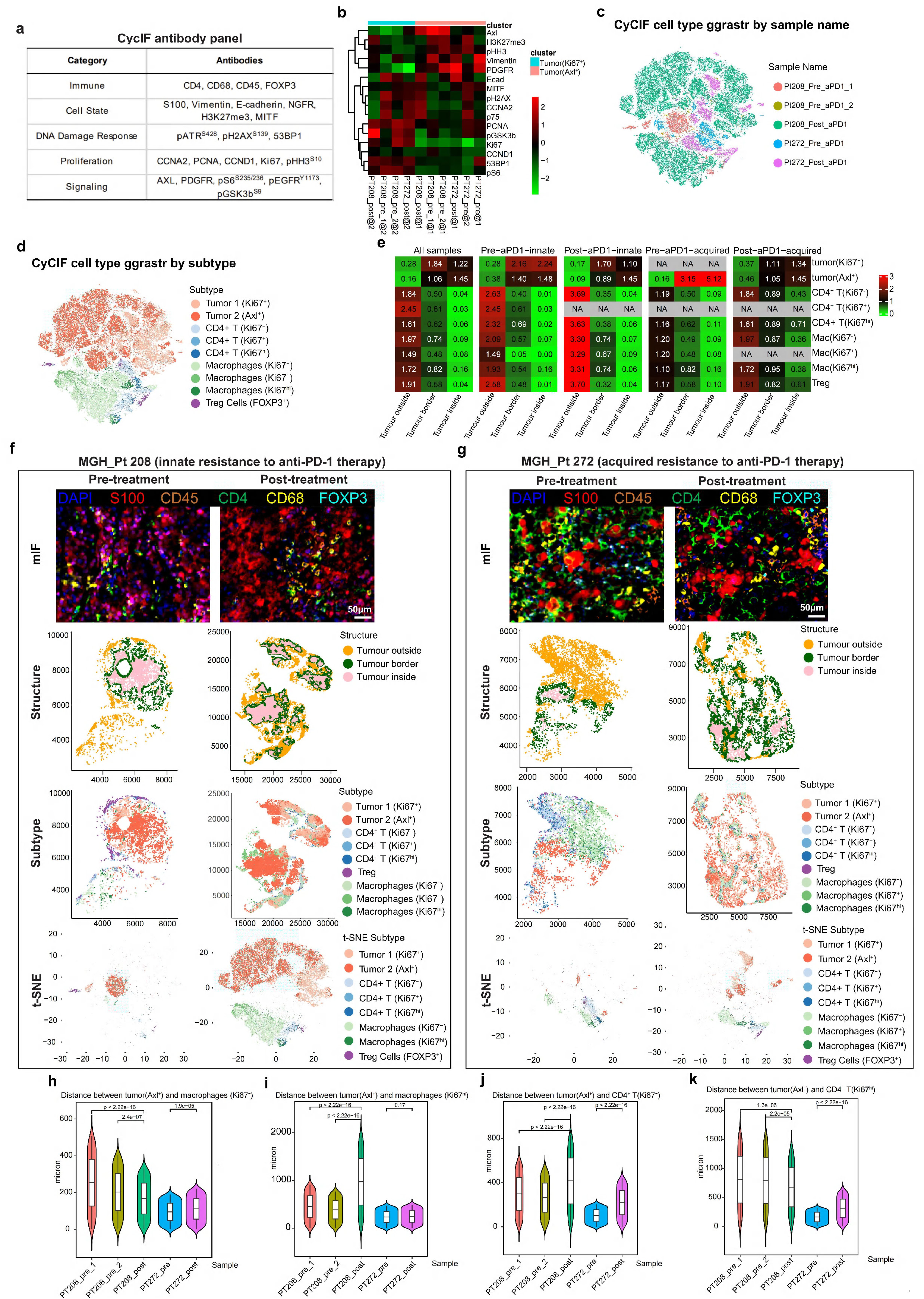
Spatially Resolved Single Cell Proteomics Analysis by CyCIF. (a) The antibody panel of 23 antibodies used in the CyCIF platform. (b) Heatmaps displaying subtypes for tumor cells, with antibodies as rows and tumor subtypes within each sample as columns. (c) t-SNE plot of cell types labeled by sample name. (d) t-SNE plot of cell types labeled by subtype. (e) The enrichment of cell subtypes within tumor outside, border, or inside. (f) Representative CyCIF figures, t-SNE plots displaying tumor structures and t-SNE plots displaying cell subtypes of tumor specimens at the time point of pre-treatment to anti-PD-1 therapy and at the time point of innate resistance to anti-PD-1 therapy derived from MGH patient 208. (g) Representative CyCIF figures, t-SNE plots displaying tumor structures and t-SNE plots displaying cell subtypes of tumor specimens at the time point of pre-treatment to anti-PD-1 therapy and at the time point of acquired resistance to anti-PD-1 therapy derived from MGH patient 272. (h) Distance between tumor (Axl ^+^) and macrophage (Ki67^−^) (micron). (i) Distance between tumor (Axl ^+^) and macrophage (Ki67^+^) (micron). (j) Distance between tumor (Axl ^+^) and CD4+ T cells (Ki67^−^) (micron). (k) Distance between tumor (Axl ^+^) and CD4+ T cells (Ki67^+^) (micron).

The clustering of CyCIF data at the single cell level revealed distinct subtypes of Ki67^+^ and AXL^+^ tumor cells (**Fig. 7b** and **Fig. S5a**). And this clustering of CyCIF data can clearly distinguish different tumor samples, cell types, and cell markers (**Fig. 7c**, **Fig. 7d** and **Fig. S5b-d**). We observed a obvious reduction in ALX^+^ tumor cells within the tumors that acquired resistance to anti-PD-1 treatment, whereas the change of AXL^+^ tumor cells was slight in tumors of innate resistance to anti-PD1 treatment (**Fig. 7e**). In addition, we observed substantial increases in percentages of CD4^+^ T cells, macrophages and Treg cells within and surrounding the tumor upon the acquired resistance to anti-PD-1 treatment occurred; however, these changes were slight in tumors of innate resistance to anti-PD1 treatment (**Fig. 7e**).

The detailed analysis of two representative patients – Pt 208, with innate resistance to anti-PD-1 therapy, and Pt 272, with acquired resistance to anti-PD-1 therapy, highlighted notable changes in cell types and states (**Fig. 7f** and **Fig. 7g**). For Pt 208, the percentages of CD4^+^ T cells, macrophages and Treg cells within and surrounding the tumor were almost unchanged between pre- and post-anti-PD1 treatment. Conversely, for Pt 272, the percentages of CD4^+^ T cells, macrophages and Treg cells within and surrounding the tumor were substantially increased after acquired resistance to anti-PD-1 treatment (**Fig. 7f** and **7g**). We also investigated the distance between tumor, and macrophages and CD4^+^ T cells, respectively (**Fig. 7h-k**). The distance between AXL^+^ tumor and Ki67^−^ macrophages, AXL^+^ tumor and Ki67^−^ CD4^+^ T cells, and AXL^+^ tumor and Ki67^+^ CD4^+^ T cells were significantly increased after acquired resistance to anti-PD1 treatment (**Fig. 7h**, **Fig. 7j** and **Fig. 7k**). However, we observed significantly decreased distance of AXL^+^ tumor and Ki67^−^ macrophages, and AXL^+^ tumor and Ki67^+^ CD4^+^ T cells (**Fig. 7h** and **Fig. 7k**), but significantly increased distance of AXL^+^ tumor and Ki67^+^ macrophages, and AXL^+^ tumor and Ki67^−^ CD4^+^ T cells after anti-PD1 treatment with innate resistance (**Fig. 7i** and **Fig. 7j**). We also focused on the Ki67^+^ CD4^+^ T cells and Ki67^+^ macrophages, and increased expression of PCNA in both Ki67^+^ CD4^+^ T cells and Ki67^+^ macrophages were observed upon the acquired resistance to anti-PD-1 treatment occurred rather than after anti-PD1 treatment with innate resistance (**Fig. S5e** and **Fig. S5f**).

Taken together, these findings underscored dynamic changes in cell types and states that were closely associated with resistance to anti-PD-1 therapy. They provided critical insights into the complex tumor immune microenvironment and highlighted potential therapeutic targets for overcoming resistance to ICB therapies.

## Discussion

The landscape of immune checkpoint blockade (ICB) therapies in metastatic melanoma has evolved significantly, yet challenges persist in understanding the molecular mechanisms governing patients’ responses and resistance. This study presents a comprehensive analysis of transcriptomic profiles from patient-matched pre- and post-treatment tumor samples derived from individuals with metastatic melanoma exhibiting either innate or acquired resistance to ICB therapies. Our findings reveal a cross-resistance meta-program of key signaling pathways that are essential for tumor progression, which is activated in metastatic melanomas with acquired resistance to ICB therapies. Conversely, a meta-program comprising crucial immune signaling pathways is suppressed in these resistant tumors. Through in-depth analysis, we identified several key pathways implicated in mediating resistance, including those associated with cell cycle and proliferation, invasiveness and metastasis, c-MYC signaling, stem cell signaling, tumor marker genes, drug resistance, gene signatures of poor survival, and Wnt/β-catenin signaling. Furthermore, spatially-resolved, image-based immune monitoring analysis by using NanoString’s DSP and CyCIF showed infiltration of suppressive immune cells in the tumor microenvironment of melanoma with resistance to ICB therapies. These findings underscore the complex interplay between tumors and the immune system, highlighting the multifaceted nature of molecular mechanisms driving resistance.

Emerging evidence have suggested that dysregulation of the cell cycle contributes as tumor-intrinsic escape mechanisms to ICB therapies^35^. The cell cycle, governed by a complex network of regulatory proteins and checkpoints, plays a crucial role in coordinating cell growth, division, and proliferation. A single-cell transcriptomic study of melanoma patients treated with ICB identified a resistance programme driven by CDK4/CDK6^36^. A previously published data sets used to study the effect of CDK4/CDK6 inhibition on breast cancer cells and mouse models, the resistance programme was repressed in response to CDK4/CDK6 inhibition. CDK4/CDK6 acts by phosphorylating the tumor suppressor retinoblastoma-associated protein 1 (RB1), and consistent with this, CDK4/CDK6 inhibition repressed the resistance programme in two RB-sufficient melanoma cell lines but not in an RB-insufficient melanoma cell line^36^. Furthermore, cell cycle-driven alterations in the tumor microenvironment, such as hypoxia, acidosis, and nutrient deprivation, can impair T cell function and infiltration, attenuating the anti-tumor immune response^37^. Targeting cell cycle dysregulation represents a promising strategy to overcome resistance to ICB therapies and enhance anti-tumor immunity. Preclinical studies have shown that pharmacological inhibitors of CDKs, such as CDK4/6 inhibitors, can synergize with ICB therapies to enhance T cell-mediated anti-tumor responses and overcome resistance in various cancer models^36,38,39^. Combination strategies targeting both cell cycle checkpoints and immune checkpoints have shown promise in preclinical studies, suggesting a feasible approach to overcoming resistance and improving outcomes for patients with metastatic melanoma.

MYC dysregulation has also been highlighted as a critical factor in mediating resistance to ICB therapies^40,41^. MYC is a transcription factor that plays a pivotal role in regulating cell growth, proliferation, and metabolism, and its aberrant expression is frequently observed in various cancer types, including melanoma^42^. MYC overexpression has been associated with the suppression of tumor-infiltrating lymphocytes (TILs), impairing the immune system’s ability to recognize and eliminate cancer cells^43^. Besides, MYC-driven tumors often exhibit metabolic reprogramming, such as increased glycolysis and glutamine metabolism, creating an immunosuppressive microenvironment that promotes immune evasion^44,45^. Given the central role of MYC dysregulation in mediating ICB resistance, targeting MYC signaling pathways represents a promising strategy to overcome resistance and enhance immunotherapy efficacy. Preclinical studies have shown that pharmacological inhibition of MYC or its downstream effectors can restore T cell-mediated antitumor immunity and sensitize to ICB therapies^46^. Additionally, combination strategies targeting both MYC and immune checkpoints have shown synergistic effects in preclinical models, suggesting a potential therapeutic approach for patients with MYC-driven cancers resistant to ICB therapies^46^.

Our study has several limitations, including the retrospective nature of the analysis, the small number of paired samples, and the inherent heterogeneity of metastatic melanoma. Future prospective studies incorporating multi-omics approaches and longitudinal sampling are warranted to validate our findings and elucidate the dynamic interplay between genomic alterations, immune cell phenotypes, and therapeutic response.

In conclusion, our study defines the transcriptomic landscape and characterized the tumor immune microenvironment associated with resistance to ICB therapies in metastatic melanoma. By delineating the molecular mechanisms underlying resistance, we aim to inform the development of novel therapeutic strategies to improve the efficacy of immunotherapy and ultimately benefit patients with advanced melanoma.

## Acknowledgements

Funding: Dr. Keith T. Flaherty was funded by Adelson Medical Research Foundation for the efforts devoted to this study. Dr. Meenhard Herlyn was funded by NIH grants P50 CA261608, P01 CA114046, U54 CA224070, and the Dr. Miriam and Sheldon G. Adelson Medical Research Foundation.

## Methods

### Patient Cohorts Treated with Immune Checkpoint Blockade (ICB)

This study incorporated data from a newly assembled cohort of 15 patients with metastatic melanoma from Massachusetts General Hospital (MGH) who were treated with immune checkpoint blockade (ICB) therapies. These data were combined with two previously published datasets (GEO accession numbers GSE115821 (https://www.ncbi.nlm.nih.gov/geo/query/acc.cgi?acc=GSE115821) and GSE168204(https://www.ncbi.nlm.nih.gov/geo/query/acc.cgi?acc=GSE168204) and a new dataset ^20,47^. Tumor specimens were collected at pre-treatment, on-treatment, and/or post-treatment stages. All patients provided informed consent, and the study adhered to relevant ethical regulations (Dana-Farber Cancer Institute IRB protocol 11-181 and The Wistar (RRID: RGD_13508588) Institute Human Subjects protocol 2802240).

We also included data from two published cohorts with bulk RNA sequencing (RNAseq) of patient-matched pre- and post-treatment tumor specimens: Lee *et al*. (EGA accession number EGAD00001005738, https://ega-archive.org/datasets/EGAD00001005738) and Johnson *et al*. (RNAseq data obtained from the corresponding author) ^18,19^. The study encompassed pre- and on-treatment samples from 47 patients with metastatic melanoma who received anti-PD-1/PD-L1 therapy. This included 42 patients from the Riaz *et al.* (GSE91061, https://www.ncbi.nlm.nih.gov/geo/query/acc.cgi?acc=GSE91061) cohort and 5 patients from the MGH cohort^25^. Additionally, data from 5 patients with recurrent glioblastoma (rGBM) who received anti-PD-1 therapy were included from Zhao *et al.* (SRA accession number PRJNA482620, https://www.ncbi.nlm.nih.gov/bioproject/482620), with bulk RNAseq data of patient-matched pre- and post-treatment tumor specimens^26^.

### Patient Cohorts Treated with Molecularly Targeted Therapies

We incorporated data from 25 patients with metastatic melanoma treated with BRAF inhibitor monotherapy who subsequently experienced relapse, derived from Kwong *et al.* (EGA accession number EGAD00001001306, https://ega-archive.org/datasets/EGAD00001001306) and Hugo *et al*. (GEO accession number GSE65185, https://www.ncbi.nlm.nih.gov/geo/query/acc.cgi?acc=GSE65185)^27,28^. Additionally, bulk tumor RNAseq data at pre- and post-BRAF inhibitor stages were included from one patient who received BRAFi at MGH (Dana-Farber Cancer Institute IRB protocol 11-181).

### RNA Sequencing and Data Processing

The RNA sequencing (RNAseq) methodology has been previously described^48^. Briefly, RNA was extracted from fresh frozen tumors using the Qiagen RNeasy Mini kit. RNA libraries were prepared with 250 ng of RNA per sample following standard Illumina protocols. RNA sequencing was performed at the Broad Institute (Illumina HiSeq2000) and the Wistar Institute (Illumina NextSeq 500). Paired fastq files were aligned to the GRCh37 reference genome using the STAR (RRID: SCR_004463) aligner with default settings^49^. Read counts were summarized using featureCounts (RRID: SCR_012919), counting only paired-ended, non-chimeric, high-quality reads (mapping quality ≥ 20)^50^. The R package edgeR (RRID: SCR_012802) was used to normalize sequencing depths and gene lengths, generating RPKMs (Reads Per Kilobase of transcript per Million mapped reads)^51^.

### NanoString nCounter® RNA and Protein Assays

The nCounter platform profiles a highly multiplexed combination of RNA and protein panels for an integrated biological view of a single sample. Two FFPE slides were used for the analysis of RNA and protein expression, respectively. Total DNA and RNA were directly extracted from FFPE slides and hybridized with specific codesets. DNA-barcode antibody panels were used for protein analysis, with barcodes obtained from UV light cleavage. After probe hybridization, RNA and protein tagsets were combined in a single hybridization reaction. Digital fluorescent barcodes were subsequently quantified for each target molecule on the nCounter platform. Data were normalized using ERCC positive loading controls and the geomean of all targets across samples, with housekeeping genes used as internal controls. SNV data were processed using nSolver 4.0 (RRID: SCR_003420). Antibodies can be found in Table S4.

### NanoString GeoMx® Digital Spatial Profiling (DSP)

The GeoMx® DSP platform allows spatially defined collection of oligonucleotide tags cleaved from specific validated antibodies on a single FFPE slide. Single FFPE slides from pre-treatment, on-treatment, and post-treatment stages of immunotherapy were selected. A multiplexed cocktail of primary antibodies with UV-light photocleavable indexing oligonucleotides and two fluorescent markers: S100B (Alex532, Cat. No. NBP2-45267AF532) and CD45 Alexa594, RRID: AB_2563458) and Syto13 (DNA staining) were applied to a slide-mounted FFPE tissue section. High-resolution images of tissue areas of interest were acquired by DSP-integrated fluorescence microscopy. Regions of interest (ROIs) for molecular profiling were selected based on dual S100/CD45 staining to optimize the assessment of immune infiltration in the tumor microenvironment. ROIs were segmented into two areas of interest (AOIs) according to the immunofluorescence intensity of S100 and CD45. Decoupled oligonucleotides from profiling reagents were rapidly aspirated and deposited into microtiter plate wells, indexed to the AOI on the tissue. After processing, indexing oligos were hybridized to NanoString optical barcodes and quantified on the nCounter platform. Data were normalized with positive controls (ERCC) and geomean across all probes. Quality control excluded AOIs with less than 30 nuclei counts.

### Cyclic Immunofluorescence (CyCIF)

Cyclic Immunofluorescence (CyCIF) is a spatially oriented single-cell proteomics approach that detects multiple proteins on a single FFPE slide. FFPE slides were treated with sequential rounds of staining using fluorescent-conjugated primary antibodies, followed by quenching with a solution of 3% peroxide and 20 mM NaOH in PBS. Images were acquired at 10X magnification using an Axioscan fluorescence slide scanner (Zeiss). Autofluorescence levels were subtracted from each image. Images were registered using Matlab (RRID: SCR_001622), and cells were segmented using QI software. Protein mean intensities and cell coordinates were analyzed using custom Python (RRID: SCR_008394) scripts available on GitHub (RRID: SCR_002630). Antibodies can be found in Table S7.

### RNA-Seq Data Batch Effect Correction

To correct for batch effects, ComBat-seq techniques were applied to raw read counts. The batch effect, a major technical variation between different datasets, was removed using the SVA R package according to the COMBAT method.

### Differentially Expressed Gene Analysis

Differential expression analysis was conducted using DESeq2 (RRID: SCR_000154), an R package. DESeq2 models gene count expression data and calculates Log2 fold change to estimate gene changes between comparison groups^52^. Two-sided Wald test statistics were computed to examine differential expression across sample groups, and p-values were corrected for multiple testing using the FDR/Benjamini-Hochberg method.

### Gene Set Enrichment Analysis (GSEA)

GSEA (RRID:SCR_003199) was implemented using the fgsea R package, which utilized pre-ranked gene lists based on differential expression results and curated gene sets (Hallmark gene set, ImmuneSigDB gene set, CGP: chemical and genetic gene set) from the Molecular Signatures Database. The permutation p-value was set at 10,000. GSEA results were visualized using volcano plots.

### Single-sample GSEA (ssGSEA)

RNA-Seq gene raw counts were normalized to transcripts per kilobase million (TPM) expression values using GENCODE (RRID: SCR_014966) version 35 as the reference transcript database. The GSVA R package computed ssGSEA values for selected pathways^53^. Heatmaps were used to visualize ssGSEA values, with hierarchical clustering performed using the “minkowski” method for distance measurement and the “ward.D” method for clustering.

In Figure 2a, ssGSEA difference values between pre-treatment and on/acquired resistance conditions were calculated for each patient. T-tests compared ssGSEA difference values between “Melanoma anti-PD-1 R Pre vs On” and “Melanoma anti-PD-1 R Pre vs Acquired Resistance,” selecting pathways with p-values less than 0.01. Heatmaps visualized ssGSEA difference values for all cohort patients, with hierarchical clustering using the “minkowski” method for distance measurement and the “ward.D” method for clustering.

### Weighted Correlation Network Analysis (WGCNA)

We employed Weighted Gene Co-expression Network Analysis (WGCNA, RRID: SCR_003302) to examine quantifiable genes and construct a gene co-expression network, utilizing the R package ‘WGCNA’^32^. To ensure consistency with scale-free attributes, we applied a scale-free R^2^ value of 0.8. In order to reduce noise and spurious correlations, we transformed the adjacency matrix into a topological overlap matrix (TOM). The network construction and module identification were based on TOM similarity. The following parameters were set: soft-threshold power (β) = 10, “cutreeDynamic” function, and a minimum module size of 20.

### Statistical Analysis

Two-sided Welch t-tests compared ssGSEA values between pre-treatment and post-treatment samples, selecting pathways with p-values less than 0.01.

### Data Availability

Source data are provided with this paper. All patient data analyzed from published papers are referenced and publicly available. The Riaz *et al.* data used in this study are available in the GEO database under accession code GSE91061. RNAseq data from Johnson *et al*. were obtained from the corresponding author. The Lee *et al.* data are available in the EGA database under accession code EGAD00001005738. The Kwong *et al.* data are available in the EGA database under accession code EGAD00001001306. The Hugo *et al*. data are available in the GEO database under accession code GSE65185. The published MGH data are available in the GEO database under accession code GSE115821 and GSE168204. Zhao *et al*. data are available in the BioProject database (RRID: SCR_004801) under accession code SRA PRJNA482620.

### Code Availability

Codes are implemented in R and are publicly available on GitHub.

