## supplementary figures for "A Comprehensive Proteogenomic and Spatial Analysis of Innate and Acquired Resistance of Metastatic Melanoma to Immune Checkpoint Blockade Therapies"

#### Supplementary Figure1

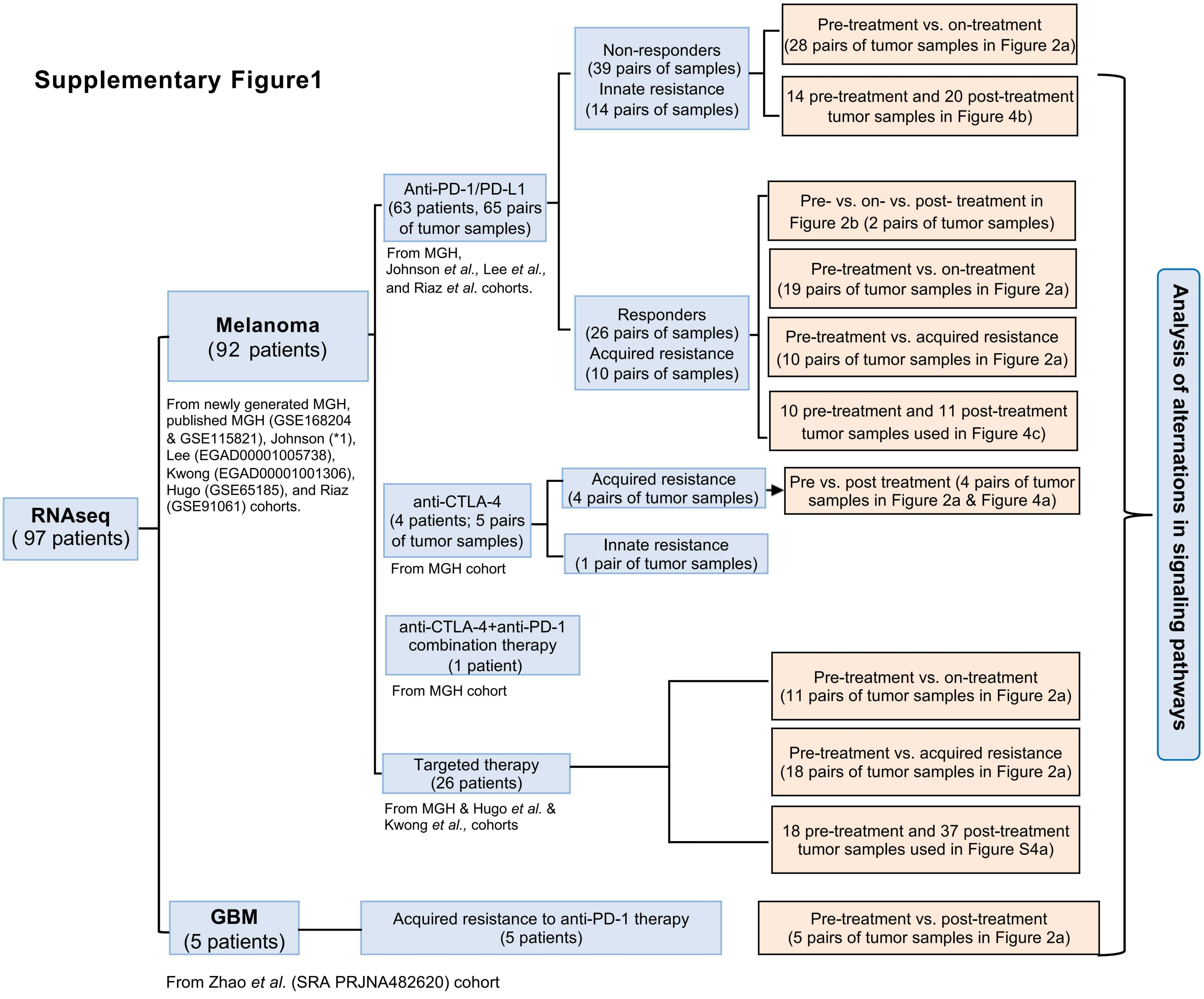


* 1: we obtained RNAseq data from the corresponding author.

### Supplementary Figure 2

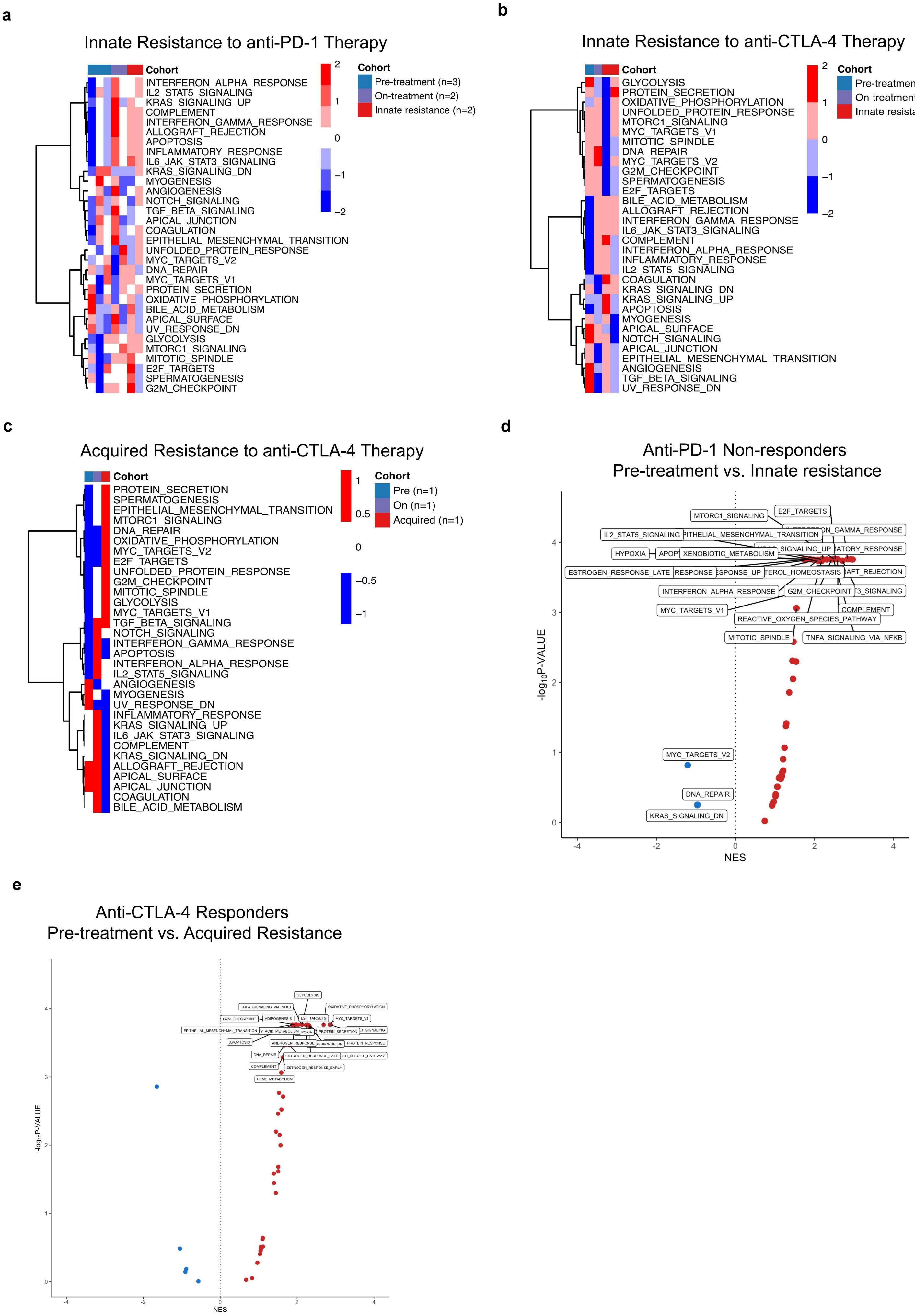

### Supplementary Figure 3

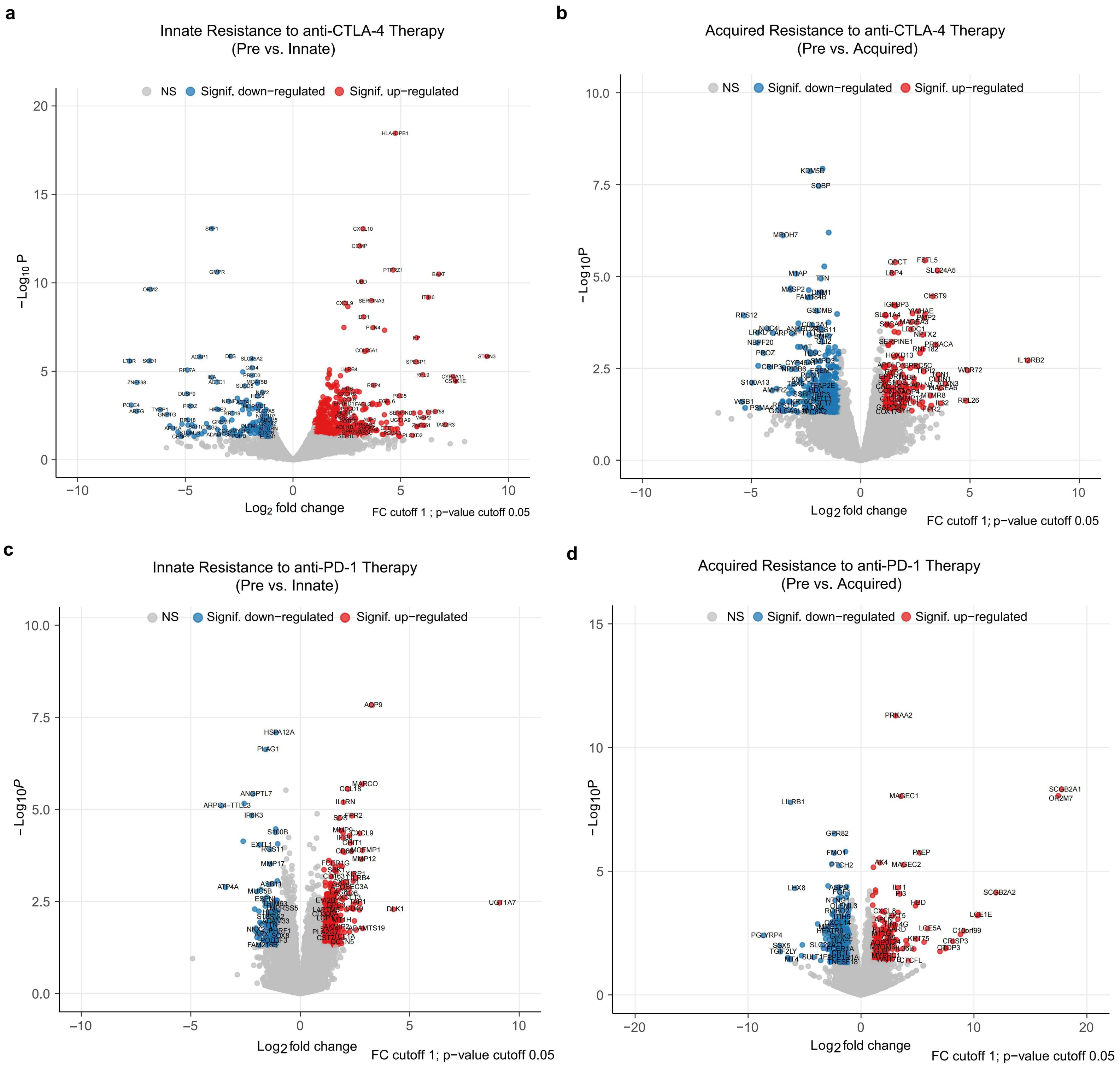

**Supplementary Figure 4**

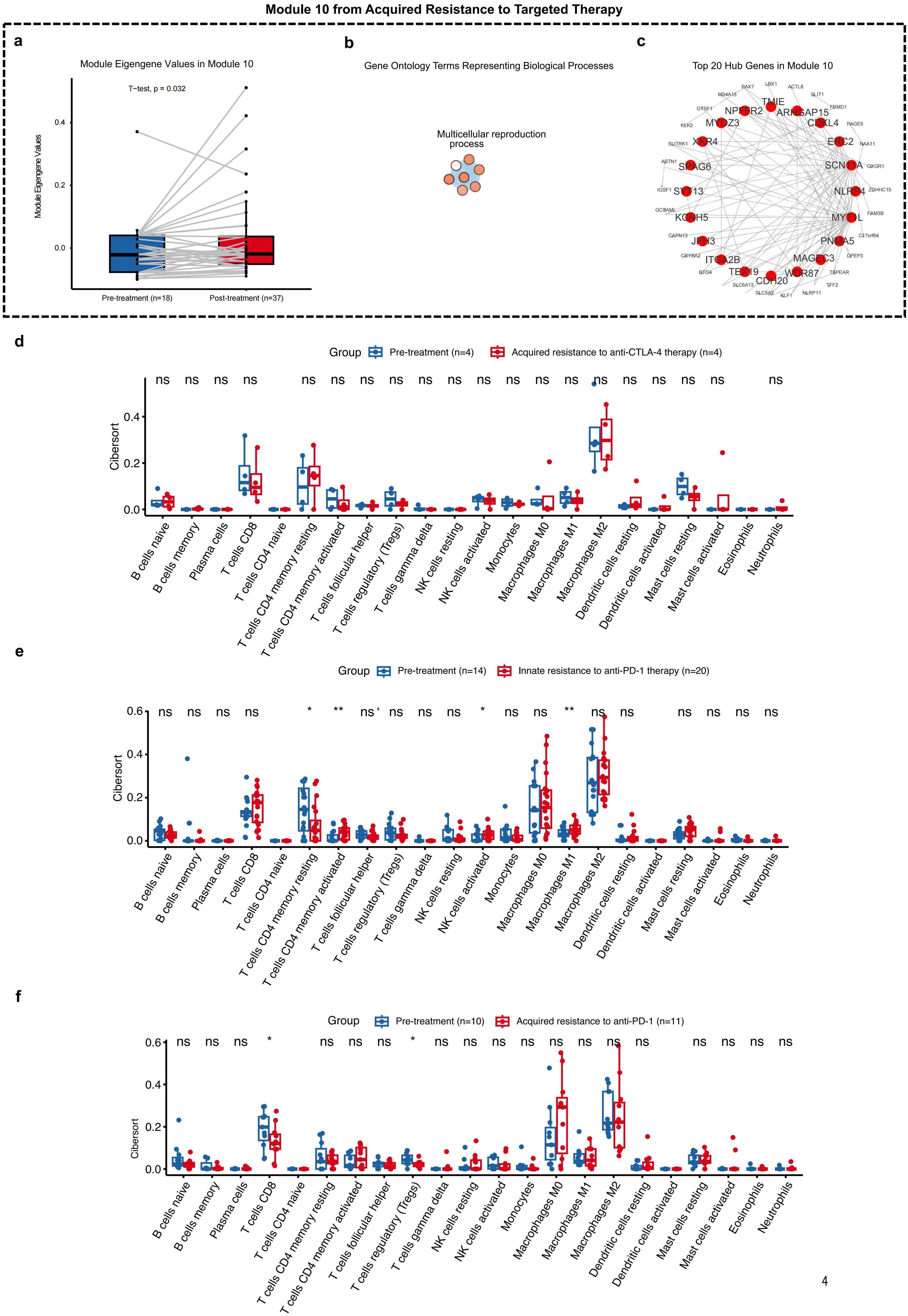

**Supplementary figure 5**

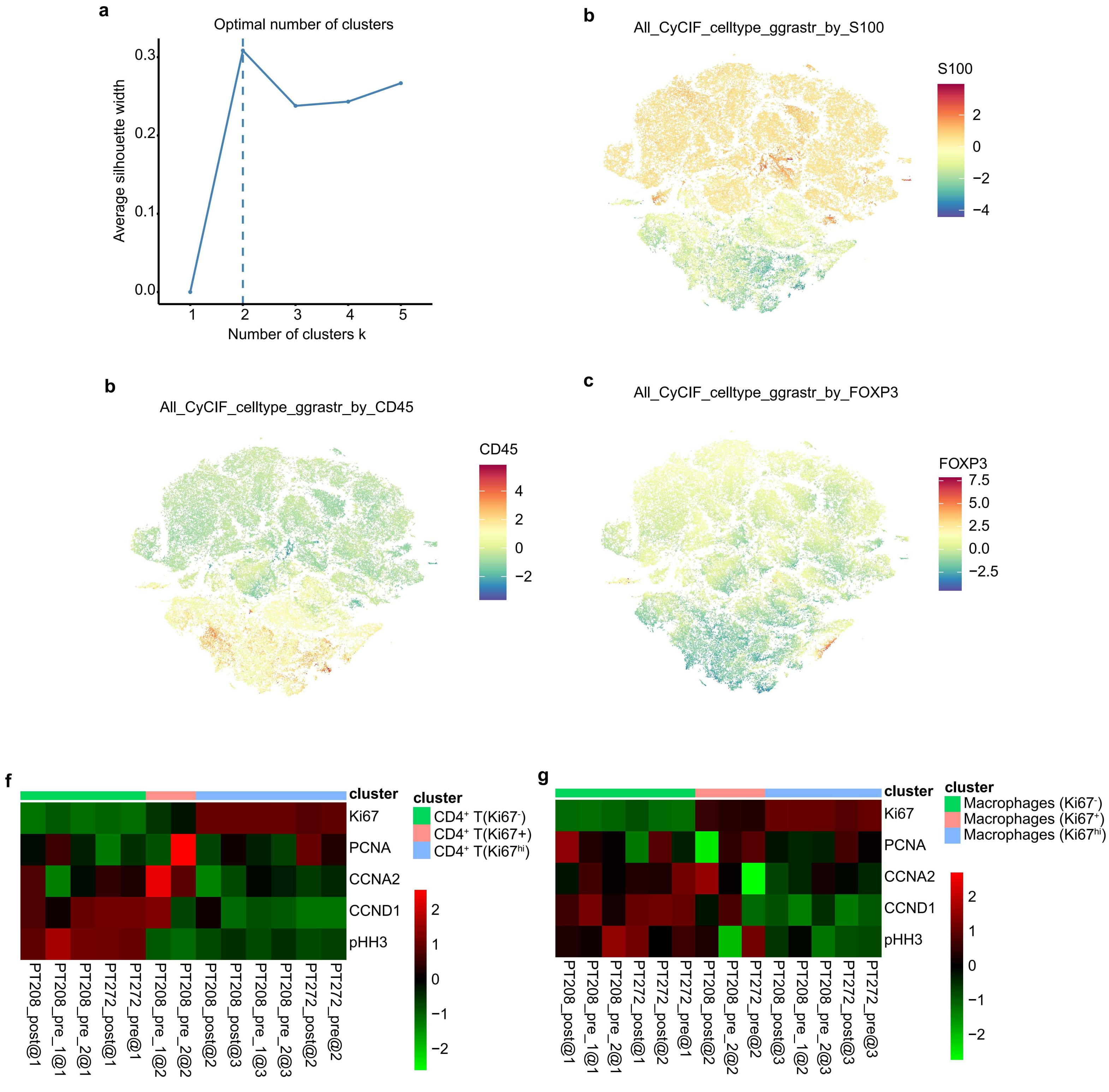
